# Single-cell immune multi-omics and repertoire analyses in pancreatic ductal adenocarcinoma reveal differential immunosuppressive mechanisms within different tumour microenvironments

**DOI:** 10.1101/2023.08.31.555730

**Authors:** Shivan Sivakumar, Ashwin Jainarayanan, Edward Arbe-Barnes, Piyush Kumar Sharma, Maire Ni Leathlobhair, Sakina Amin, Lara Heij, Samarth Hegde, Assaf Magen, Felicia Tucci, Bo Sun, Shihong Wu, Nithishwer Mouroug Anand, Hubert Slawinski, Santiago Revale, Isar Nassiri, Jonathon Webber, Adam Frampton, Georg Wiltberger, Ulf Neumann, Philip Charlton, Laura Spiers, Tim Elliott, Pallavur V. Sivakumar, Alexander V. Ratushny, Mark Middleton, Dimitra Peppa, Benjamin Fairfax, Miriam Merad, Michael L. Dustin, Enas Abu-Shah, Rachael Bashford-Rogers

**Author notes:** **Corresponding authors:** Shivan Sivakumar, Enas Abu-Shah and Rachael Bashford-Rogers. denotes joint first authors (Shivan Sivakumar, Ashwin Jainarayanan, Edward Arbe-Barnes). denotes joint second authors (Piyush Kumar Sharma, Maire Ni Leathlobhair, Sakina Amin). denotes joint last authors (Michael L. Dustin, Enas Abu-Shah, Rachael Bashford-Rogers).

## Abstract

Pancreatic ductal adenocarcinoma (PDAC) has an extremely poor prognosis. Understanding the multiple mechanisms by which the tumour evades immune control, and how these mechanisms may be disrupted is critical to developing targeted immunotherapies. Previous studies have shown that higher lymphocyte infiltration is associated with better survival, and here we investigated what mediates these differences. We performed a comprehensive analysis of PDAC-associated immune cells using single cell multi-omics coupled with re-analysis of public PDAC scRNA-seq datasets. We introduce novel single-cell and repertoire analyses that have uncoupled diverse roles and contributions of various immune cell populations within different tumour microenvironments (TMEs). They revealed clear distinctions in the clonal characteristics among different patient groups, provided valuable insights into the mechanisms of immune cell migration and tissue adaptation underlying these disparities. These results point to differential CD4 polarisation of intra-tumoural T cells, differential B cell differentiation, GC reactions, antigen presentation pathways, and distinct cell-cell communication between the myeloid-enriched and adaptive-enriched groups. Overall, we identified two major distinct themes for future immune intervention within PDAC patients between those with higher adaptive versus myeloid immune cell infiltration.

## Introduction

Pancreatic ductal adenocarcinoma has the worst survival of any common human cancer, with a 5-year survival of below 10%^1^. The mainstay of treatment is chemotherapy, however, approximately 15% of patients benefit from surgical resection, which can potentially provide cure in a subset of those patients. Despite the introduction of immunotherapy, the benefit in PDAC is minimal^2–6^, and so there is an unmet need to develop better treatments for patient benefit.

Previous work from our group and others has suggested that there is a sizable immune infiltrate in these tumours and understanding the nature of this infiltrate is critical for developing pragmatic immunotherapy strategies for PDAC^7–10^. We have previously shown that patients with high tumour lymphocyte infiltration at resection have a better prognosis than those that do not^8^. Furthermore, after characterising tumour infiltrating lymphocytes (TILs) in PDAC, we see that even though there is limited exhaustion in a subset of CD8 T cells, we observed that a significant number of CD4 and CD8 T cells were senescent^7,8^. Additionally, we see a activated Treg expressing checkpoints TIGIT, ICOS, CTLA4 and CD39^7,9^ suggesting a strongly immunosuppressive microenvironment. This activation was determined by the high expression of the checkpoints TIGIT, ICOS, CTLA4 and CD39^7,9^.

Other groups have made recent observations regarding the intra-tumoural immune infiltrate^11^. Peng et al. did the first large scale single cell analysis of PDAC and demonstrated that there was a complex immune infiltrate, and they highlighted that T cells were the dominant immune cell in the TME^10^. Steele et al. performed a second large single cell experiment and demonstrated that the predominant CD8 T cell exhaustion marker was TIGIT^12^. Schlack et al, have performed single cell sequencing with TCR sequencing. They have identified a heterogeneous lymphocyte infiltrate and trajectory analysis demonstrated similarities between inhibitory and dysfunctional populations^13^. Brouwer et al had undertaken a single cell CyTOF analysis using a 41-marker panel focused on infiltrating lymphocytes. They found low levels of tissue resident cytotoxic CD8 T cells and they concurrently have low levels of PD1. Interestingly, the group has also found high levels of activated Tregs and B cells^14^. Liudahl et al. used an immune focused multiplex IHC panel to evaluate leukocyte populations in a cohort of 135 PDAC patient samples. They demonstrated that the T cell to CD68 ratio is important in the treatment naive setting to demonstrate prognostic benefit^15^.

There is a growing body of evidence describing a distinction between lymphocyte- and myeloid-enriched tumours and understanding what is driving this is critical to therapeutic interventions. Despite the growing number of datasets aiming at defining the nature of the different immune subsets within PDAC TME, we still lack an understanding of the clonal evolution and differentiation pathways driving these populations. PDAC has been traditionally considered to have a low mutation rate, suggesting a low prevalence of antigens to stimulate the immune response. However, seeing the presence of activated and exhausted cells within the TME, and associations between cytotoxic CD8 T cells, B cells and neoantigen quality with patient survival^16,17^ suggests the presence of specific stimuli and warrants the investigation of the clonal distribution and evolution of both T and B cells. Multiple previous studies have shown that higher adaptive immune cell infiltration is associated with marginally better survival^8,18^, however, both adaptive and myeloid enriched PDAC patients have dismal prognosis^6^. This study’s objective is to elucidate the distinctive features of adaptive immune responses in patients’ tumours with high levels of adaptive and myeloid cell populations and to identify the specific immune suppression pathways that set apart the myeloid-high and adaptive-high patient groups. These insights will help in patient stratification and the development of personalised therapeutic approaches. Furthermore, the nature of B and T cells moving between the tumour and draining lymph nodes is important for mounting effective anti-tumoural immune responses and establishing long-term systemic memory^19^. However, the signals responsible for B and T cell tumour infiltration, retention and egress, such as adhesion and chemokine milieu, are unknown. We aimed to explore the nature and determinants of B and T cell immunosurveillance in PDAC to identify pathways that can be targeted to improve immune cell trafficking.

To this end, we performed the largest and most comprehensive analysis of PDAC-associated lymphocytes from tumour and blood to date using single cell multi-omics analysis coupled with the re-analysis of public PDAC scRNA-seq datasets^10,12^. Importantly, we developed and applied novel single cell analyses to uncouple the distinct roles and contributions different immune cell populations, the clonal nature across patient groups, the nature of immune cell migration and tissue adaptation, and provided insights into key pathways defining these differences. This study lays the foundation for understanding why immunotherapy has so far not been successful in PDAC, and provides an avenue for identifying novel therapeutic targets based on an enhanced understanding of the patients’ intra-tumoural immune composition.

## Results

### Single cell profiling of PDAC tumour immune cell infiltration across three datasets

To elucidate the heterogeneity of tumour immune cell infiltration, we performed single cell RNAseq (GEX), ADT-seq (cell surface protein expression derived from Antibody-Derived Tags), and BCR/TCR-seq on CD45+ cells enriched from matched PBMCs and fresh tumour tissue following surgical resection of 12 treatment-naïve patients, herein referred to as PancrImmune. In addition to the PancrImmune dataset, we integrated and re-analysed the two largest existing PDAC scRNA-seq datasets from Peng^10^ and Steele^12^ (**Figure 1a**). Integrative multi-omics analysis of GEX, ADT-seq and BCR/TCR-seq allowed for high confidence and quality annotations of B cell, T cell and myeloid populations (**Figure 1b, Supplemental Table 1**), making this the largest single cell analysis of immune cells in PDAC with robust detection of immune cell diversity, whilst also being the first PDAC study to incorporate all four single cell modalities (GEX, ADT-seq, BCR-seq and TCR-seq). This high granularity analysis of immune cells in the blood and PDAC tumour infiltrate revealed high complexity of immune cell infiltration in the TME with a wide variety of activated and regulatory immune cells in all major immune subsets (**Supplemental Item 1**). The integrated Peng and Steele datasets only had GEX data and therefore we used a novel support vector machine (SVM) cell label transfer method, *SVMCellTransfer*, using the PancrImmune GEX data as a reference (**Supplemental Item 1, Supplemental Table 1**, see methods). The resulting gene expression patterns of each cell annotation type reflected well the patterns observed in the PancrImmune reference dataset (**Supplemental Item 1**).

**Figure 1:**
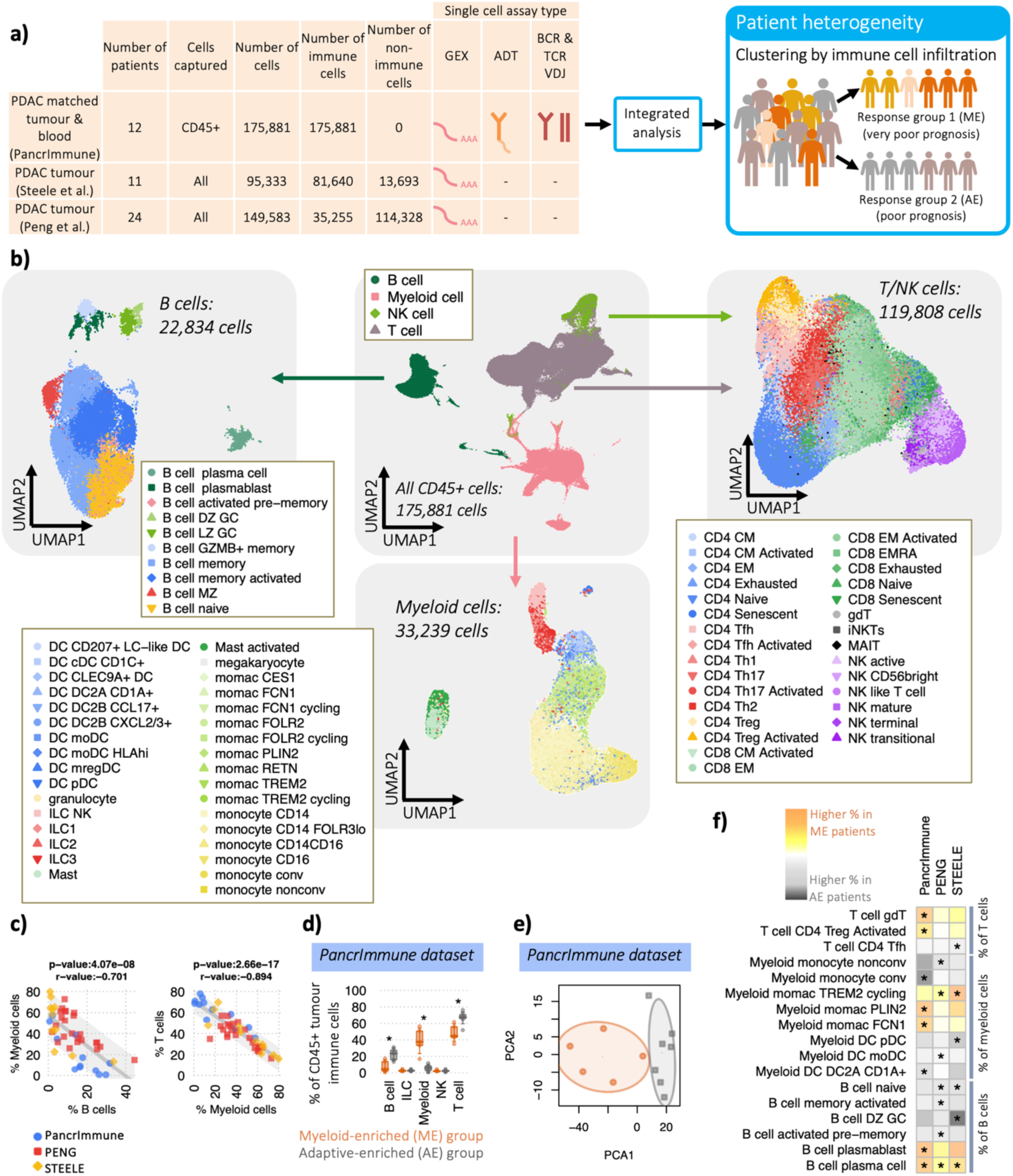
Increased intra-tumoural lymphocyte infiltration is associated with distinct immune cellular compositions. a) Schematic of the datasets and analyses b) UMAP dimensionality reduction of the immune cells from the PancrImmune dataset depicting total immune cells (centre), B cells (left), myeloid cells (bottom) and T cells (right). c) The correlation of (left) B cells and Myeloid cells and (right) myeloid cells and T cells as a proportion of total intra-tumoural immune cells across the three datasets, coloured blue red and yellow for the PancrImmune, Peng and Steele datasets respectively. d) Principal component analysis (PCA) of the immune cell proportions per sample, coloured orange for myeloid-enriched (ME) patient samples and grey for adaptive immune cell enriched (AE) patient samples (PancrImmune dataset). e) The cellular proportions of the broad immune cell types between myeloid- and adaptive-enriched patients. f) Heatmap of the differences in cellular proportions between ME and AE patient tumour samples. The colour denotes the proportional skew between ME and AE patients, and * denotes a significant difference between ME and AE patients (p-value <0.05). Statistical tests were performed by MANOVA.

### PDAC tumour myeloid infiltration positively associates with plasma cell abundance

Patients’ tumours span a spectrum of immune cell infiltration, and higher intra-tumoural T cell/lymphocyte frequencies are typically associated with improved patient survival (**Supplemental Table 2**). We therefore investigated next what might be mediating these differences. The proportion of immune cells consisting of tumour infiltrating myeloid cells inversely correlates with B and T cells consistently across datasets (p-values<1e-7, **Figure 1c**, p-values per dataset <0.018, **Table S3**). To better understand the mechanisms underlying this, we compared patients with high B and T cell tumour infiltrate proportions (as a percentage of CD45+ cells), termed adaptive-enriched (AE) versus high myeloid, low B and T cell proportions, termed myeloid-enriched (ME) (**Figure 1d-e**). This mirrors the prognostic signatures previously identified^8^ and summarised in **Supplemental Table 2**. Indeed, the top 10 differentially expressed genes (DEGs) between groups (pseudobulk analysis) have predominantly B cell and T cell specificities for AE patients, or myeloid cell specificities for ME patients (**Supplemental Table 4**). The cellular subset proportions within B cells, T cells, NK cells and ILCs significantly differed between AE and ME groups across all three datasets, and were clearly separable by PCA analysis (**Figure 1e, Supplemental Figure 1a-c**). Across all three datasets, we observe significantly reduced plasma cell abundance with increased overall B and T cell infiltration, along with consistent trends in proportions of other immune cell populations demonstrated across datasets (**Figure 1f, Supplemental Figure 1d-e**), suggesting that different mechanisms of differentiation, proliferation and recruitment may be acting in the different patient groups. Indeed, this is in agreement with previous studies showing an association between plasma cells and myeloid cells in lymphoid tissues^20–22^.

We also confirmed that there are consistent significant differences in myeloid versus adaptive immune cell infiltration into the tumour when considering the total cell population (rather than just the proportion of CD45+ cells) in both the Peng and Steele datasets (**Supplemental Figure 1f**). There were no significant differences in the proportions of non-immune cell types between the ME and AE patients, suggesting that tumour load (found in the epithelial cell compartment) and non-immune cell composition is not driving these differences. No correlations were observed with other patient factors including age, gender, prior disease history.

### Inverse correlation between ductal and immune cell subset proportions

Using the Peng and Steele datasets where all pancreatic cells were present, we were able to dissect the correlations between the immune and non-immune cell compartments. Although there was no correlation between the epithelial cells and any other non-immune cell type, we saw the strongest significant inverse correlations between epithelial (containing the tumour cells) with the myeloid, T cells and NK cell proportions (**Supplemental Figure 1f**), suggesting direct immunosuppressive activity by the tumour cells. Moreover, there was no significant inverse correlation between the fibroblast compartment and the immune cell infiltration, which supports recent evidence disputing the previously held idea that the desmoplastic core limits immune infiltration^23^.

### B cell selection is distinct between patient groups

Next, we examined the intra-tumoural B cells in greater detail given their divergent proportions between patient groups. The proportions of both plasma cells (PCs) and plasmablasts (PBs) of total B cells were significantly higher in ME tumours than AE tumours (**Figure 2a)**. IGHV gene usages and isotype features were distinct and clearly separable by PCA between AE and ME patients (**Figure 2b, Supplemental Figure 2a**, p-values<0.0125) suggesting different B cell repertoire selection processes between patient groups^24^.

**Figure 2:**
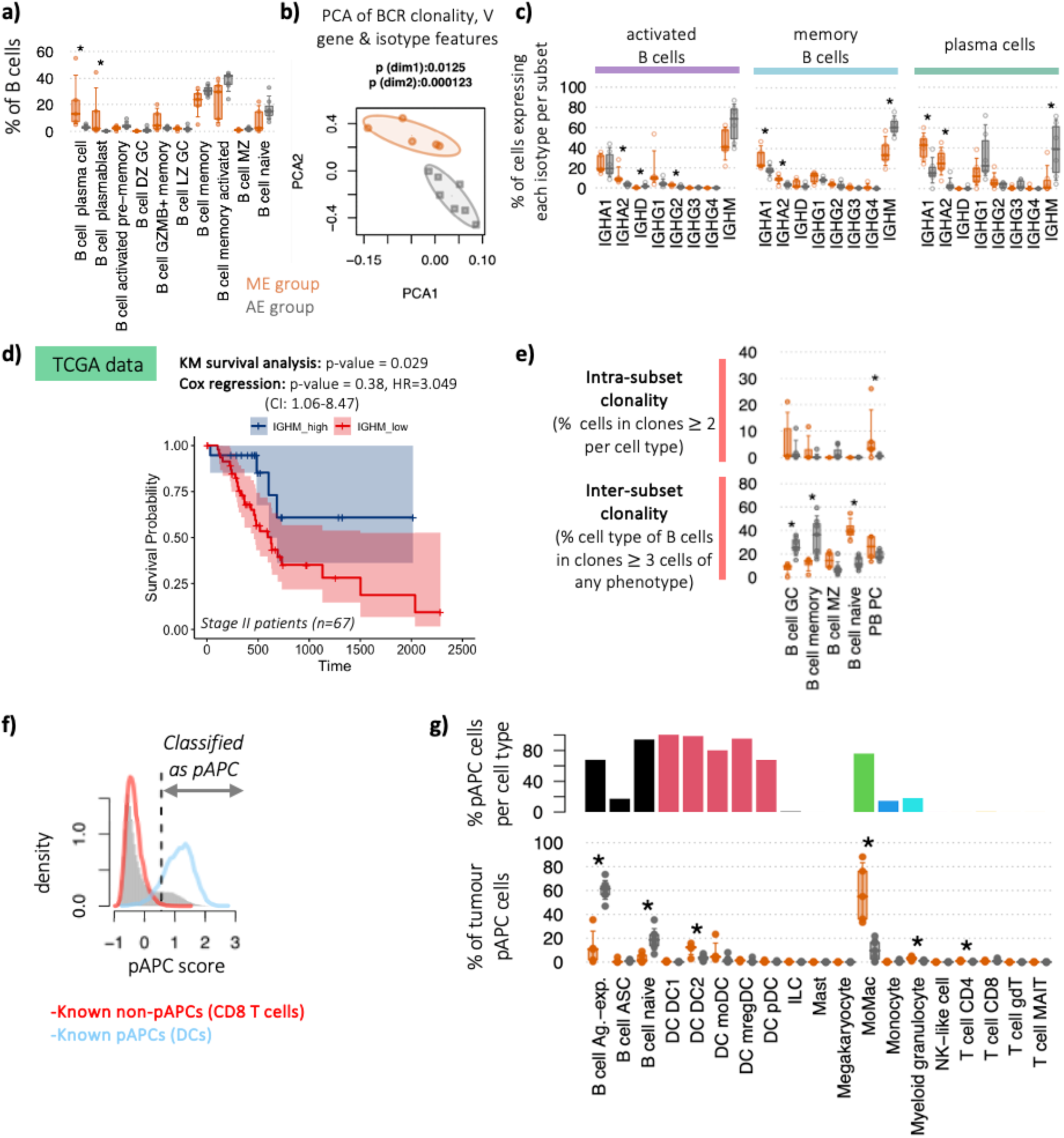
Increased PDAC lymphocyte infiltration is associated with differences in B cell selection, clonal expansion and class-switch recombination. a) Immune cell subset proportions between ME and AE patient groups within tumour B cell subsets as a proportion of total B cells (orange represents ME patients and grey represents AE patients) within the PancrImmune dataset. b) Principal component analysis (PCA) of the tumour BCR clonality, IGHV gene usages and isotype usages, coloured by patient group. c) The proportions of tumour B cells within activated, memory and plasma cells expressing each isotype, coloured by patient group. d) Clonality of the tumour B cell subpopulations between the ME and AE patient groups via two measures: (top) *intra-subset clonality* (the percentage of cells in clones >2 cells per subset, measuring the clonality within the subset thus reflecting specific cell populations which are actively expanding), and (bottom) *inter-subset clonality* (the percentage of cells of each cell type as members of clones >3 cells across all populations, demonstrating, this indicates cells within each B cell subset that may be members of larger clones than span multiple phenotypes, reflecting B cell plasticity of expanding clones). e) Survival plot for high vs low IGHM expression with a p-value for Kaplan–Meier (KM) plot (log-rank test) and Cox proportional hazards model (Wald test). HR = hazard ratio, CI = confidence interval. f) Histogram of the professional antigen presentation (pAPC) scores for (grey) all tumour cells, (red) tumour CD8 T cells and (blue) tumour DCs. Dashed line indicates the threshold for classification of pAPCs. g) (top) Bar chart of the percentages of pAPCs comprising each cell type, and (bottom) the proportion of tumour pAPCs comprising each cell type between patient groups. All analyses in this figure were performed on the PancrImmune dataset. * denotes p-values<0.05, and tests were performed by MANOVA.

### Reduced B cell class-switching in AE patients

Elevated IgA1 and IgA2 were observed in the intra-tumoural activated, memory, and antibody secreting PB cells in the ME patient group, whereas elevated IgG1 and IgM levels were observed in the AE group (**Figure 2c**). These differential phenotypes are suggestive of differences in B cell signalling and germinal centre (GC) or tertiary lymphoid structure (TLS) responses. Interestingly, the patients associated with better prognosis (AE) had reduced class switch recombination (increased proportion of IgM B cells) compared to ME. Indeed, this observation of elevated IgM in the better-prognostic patients is supported when examining the larger PDAC TCGA dataset (n=67 patients at stage II), in which we identified a weak but significant association between both high IGHM with improved survival (**Figure 2d**). In addition, IgM+ BCRs had lower levels of somatic hypermutation (SHM) (**Supplemental Figure 2b**). Taken together the AE patients may have distinctive GC reaction outcomes. We note that the dominant IGHA1, IGHA2 and IGHM isotype usages observed here also reflect what is seen in healthy pancreatic tissue (reanalysis of GTEx RNA-seq data, **Supplemental Figure 2c**). This could be suggestive that the pancreatic environment and supporting draining lymph nodes preferentially support class-switching to IGHA1/2 as observed in other GI tract locations, rather than a predominance of IGHG1/2 observed in non-GI tract tissues^25^. These differences in isotype usages were not observed in blood (**Supplemental Figure 2d**), in keeping with tissue-specific differences rather than differences in systemic B cell responses between patients. Fc receptors for IGHA (FCAR/CD89), which are known to have dual effect, either to inhibit or activate macrophage responses depending on either monovalent or multimeric IgA ligation^26^, are predominantly expressed in pancreatic myeloid cells, and Fc receptors for IGHM (FCMR) are predominantly expressed in B, T and NK cells (**Supplemental Figure 2e**). Together, this further strengthens the relationship between IGHM and improved survival (potentially as antigen presenting cells) as seen in lung cancer^27^, despite the signs of reduced GC efficiency, as seen in the AE group, and potentially the relationship between IGHA secretion and myeloid cell phenotypes driving one of the pathways of immune suppression in the ME patients.

### Increased GC B cell clonality in AE patients

We next assessed the clonality of the B cell subpopulations via two measures: intra-subset clonality which reflects specific cell populations which are actively expanding, and inter-subset clonality to reflect the expansion and differentiation between subpopulations (**Figure 2e**). Intra-subset clonality quantifies the percentage of cells in clones of 2 or more cells per subset, measuring the clonality within the subset. Inter-subset clonality quantifies the percentage of cells of each cell type as members of clones of 3 or more cells across all populations, this indicates cells within each B cell subset that may be members of larger clones that span multiple phenotypes, reflecting B cell plasticity driven by the specific TME signals they encounter. The elevated clonality observed in ME of antibody secreting cells (PCs and PBs) suggests that these are arising from recent or ongoing immune reactions in ME patients. The relatively higher levels of inter-subset clonality in the naïve B cells in ME patients, despite the expectedly low intra-subset clonality, is likely driven by the activation and clonal expansion of some naïve B cells and transition to other B cell phenotypes. The highest intra-subset clonality was observed in the GC B cells in keeping with these cells partaking in clonal B cell response, potentially in TLSs. Indeed, these comprised the largest proportion of B cells from expanded clones (inter-subset clonality) along with memory B cells in the AE group only, with significantly lower inter-subset clonality in these cells in the ME patients. There were no significant differences in the proportions of GC B cells, which may be due to the small numbers of patients and GC cells. Together, this is suggestive of GC formation in AE patients with greater clonal expansion in GC B cells, however, resulting in unswitched memory B cells rather than intra-tumoural plasma cells. Conversely, GC B cells in ME patients are not as clonal, however, the responses are predominantly IGHA1 and IGHA2, and are more likely to differentiate into plasma cells.

### B cells comprise a major pool of antigen presenting cells in AE patients

B cells are considered as one of the major professional antigen-presenting cells (pAPCs) via the MHC II pathway^28^, however their role in the activation of T cells in PDAC has not been fully explored. Here, we derived a pAPC score for each cell by quantifying the feature expression programme for MHC II and accessory pathway molecules (see **Methods**). We defined pAPCs as those above a threshold derived from the optimal separation between scores from DCs (known pAPCs) and CD8 T cells (known non-pAPCs) (**Figure 2f**). Together with DCs, >65% of naive and antigen-experienced (activated and memory) B cells and monocyte-derived macrophages (MoMacs) are pAPCs (**Figure 2g**). Whilst a mean of 57.6% of pAPC are MoMacs, and 21.1% of pAPC are DCs in the ME patients, only 16.0% of pAPCs are B cells (**Figure 2g**). Interestingly B cells comprise a major source of antigen presenting cells in the AE patient group, 80.4% of pAPCs are B cells (p-values<0.05), mainly antigen-experienced (activated and memory) B cells. This trend is validated in the Peng et al dataset (**Supplemental Figure 2f-i**, p-values<0.05). Given the elevated T cell infiltration in the AE patients (**Figure 1f**) and the significant contribution of B cells to the pAPC pool, this highlights a potential role for B cells in PDAC TME in shaping T cell activation.

### Increased CD8 T cell clonality in AE patients

To understand the drivers behind the increased T cell prevalence in the AE patients, we performed clonality analysis of the T cell populations using the TCR sequencing data. We observe an increased CD8 T cell clonality in the AE group (**Figure 3a**, p-values<0.05) which suggests that the increased PDAC lymphocyte infiltration is partly driven by clonal activation and expansion. This is observed in both tumour-infiltrating and peripheral blood T cells, implying that some of the tumour expanded clones could potentially be also present in blood. We further analysed the different sub-populations for any differential presence of T cell subsets between the two groups of patients. In addition to the elevated CD4:CD8 T cell ratio in ME patients (**Supplemental Figure 1a**, p-value<0.05), ME tumours showed higher proportions of Treg, activated Treg, and gamma-delta (gd) T cells, while AE tumours had higher proportions of T follicular helper (fh), naïve CD4 cells and CD8 effector memory (EM) cells (**Figure 3b**). The increased proportion of CD8 EM cells in the AE group further reinforces the role of clonal activation in driving T cell infiltration, and the elevated proportions of Tfh cells in the AE patients further supports the inter-relationship between B and T cells in PDAC. By contrast, the elevated proportions of Tregs and activated Tregs in the ME group could reflect the increased immunosuppressive TME of these tumours. A validation was performed via cellular deconvolution of the PAAD TCGA dataset (n=156 patients) showing that indeed Treg proportions (as a proportion of total T cells) correlated with myeloid cell proportions (as a proportion of total immune cells) (**Supplemental Figure 3a**), whereas the T cell proportion of total immune cells inversely correlated with the proportion of myeloid cells (**Supplemental Figure 3a**). Furthermore, TCR clonality and TRVB features are distinct between AE and ME patients (**Supplemental Figure 3b**: PCA plot of T cell repertoire features, **Supplemental Figure 3c**: V gene usages). Together, this is suggestive of different T cell selection processes between ME and AE patients.

**Figure 3:**
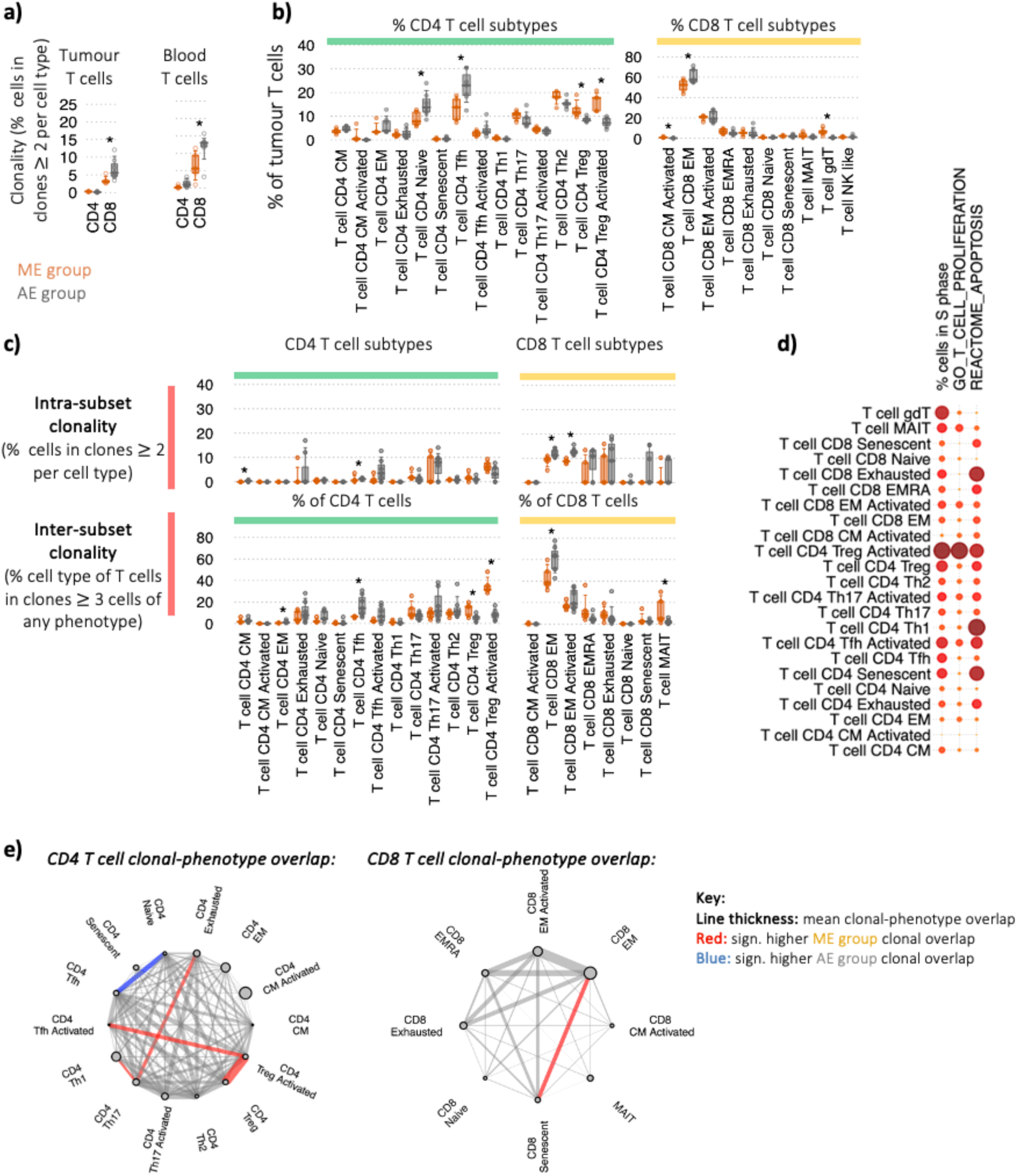
Increased PDAC lymphocyte infiltration is associated with increased activated Treg clonality and proliferation. a) The clonality of tumour T cells within total CD4 and CD8 T cell populations, measured by percentage of clones consisting of >2 cells. Orange represents ME patients and grey represents AE patients. b) Immune cell subset proportions between ME and AE patient groups within tumour T cell subsets as a proportion of total CD4 and CD8 T cells, respectively. c) Clonality of the tumour T cell subpopulations between the ME and AE patient groups via two measures: (top) *intra-subset clonality* (the percentage of cells in clones >2 cells per subset, measuring the clonality within the subset thus reflecting specific cell populations which are actively expanding), and (bottom) *inter-subset clonality* (the percentage of cells of each cell type as members of clones >3 cells across all populations, demonstrating, this indicates cells within each T cell subset that may be members of larger clones than span multiple phenotypes, reflecting T cell plasticity of expanding clones). d) The relative mean percentage cells within predicted S phase, mean Gene Ontology (GO) T cell proliferation score, and mean REACTOME apoptosis scores per cell between tumour T cell populations. The circle size indicates the relative means between cell types. e) Level of tumour TCR clonal sharing between (left) CD4 T cell and (right) CD8 T cell populations. Each line represents a sharing of TCR clones between cell types, and the line thickness denotes the mean relative level of sharing. A red line denotes that the clonal sharing between the corresponding cell types is significantly higher in the ME patients than AE, and a blue line denotes that the clonal sharing between the corresponding cell types is significantly lower in the ME patients than AE. The size of the dot represents the mean relative frequency of the corresponding cell type. All analyses in this figure were performed on the PancrImmune dataset. * denotes p-values<0.05, and tests were performed by MANOVA.

### Activated Tregs are enriched in expanded T cell clones and are the most proliferative T cell population

We next assessed the clonality of the T cell subpopulations via intra-subset clonality and inter-subset clonality (**Figure 3c**). Intra-subset clonality measures the clonality within the subset, and inter-subset clonality quantifies the cells within each T cell subset that may be members of larger clones that span multiple phenotypes, reflecting T cell plasticity driven by the specific TME signals they encounter. The highest intra-subset clonality was observed in the CD8 EM, activated EM, effector memory cells re-expressing CD45RA (EMRA), exhausted and senescent T cells. There was preferential intra-subset expansion in the AE group of the CD8 EM and CD8 activated EM T cells. Taken with the increased percentages we observed earlier (**Figure 3b**), this provides additional support that the increased CD8 EM presence in AE patients is driven by local expansion within the tumour. Notably, although activated Tregs, which are marked by the highest expression of immunomodulatory molecules TIGIT, ICOS, and CTLA4, as well as the transcription factors FOXP3 and IKZF2, were not the most clonal CD4 T cell population (intra-subset clonality), which is to be expected from a polyclonal regulatory T cell population, they were members of the most expanded clones (inter-subset clonality, **Figure 3c**), which was significantly more expanded in the ME patient group, suggesting some of the observed Tregs could be differentiating from other CD4 T cells within the TME.

The factors driving CD4 T cell differentiation toward CD4 Treg phenotype in PDAC are not yet understood, and whether increased proliferation or reduced apoptosis in the Treg populations were causing this association with clonal expansion. Indeed, activated Tregs represent the most proliferative lymphocyte population within the tumour (as measured by percentage cells in S phase and mean per cell GO_T_CELL_PROLIFERATION score across all patients, **Figure 3d**). Conversely, there was no notable difference in apoptosis, as measured by mean per cell REACTOME_APOPTOSIS pathway score. There were no widespread significant differences between patient groups. This is suggestive that activated Tregs are associated with clonal expansions and are the most proliferative T cell population within the tumour.

### Distinct T cell clonal fate between AE and ME patient groups

Through quantifying the relative overlap of clones between different phenotypes within the CD4 and CD8 T cell populations, lineage patterns can be discerned (**Figure 3e**). Indeed, highest clonal overlap in the CD8 T cell populations was observed between CD8 EM, activated EM and CD8 senescent T cell subsets, suggesting that common antigens are driving the expansions across these populations. Elevated intra-subset clonality was observed in the ME patients between CD8 EM and CD8 senescent T cell populations in the tumour, suggesting that activated T cells are pushed to dysfunctional phenotypes. The development of senescence in both patient groups suggest that the TME is conducive to the generation of these populations through potentially shared pathways. In the CD4 T cells, activated Tregs have the highest degree of clonal overlap with activated Tfh and activated Treg populations, which was significantly higher in the ME patients. In AE tumours there was higher clonal overlap between CD4 naïve and Tfh T cells. Taken together, these results point to differential CD4 polarisation of intra-tumoural CD4 T cells between the ME and AE groups.

### T cell clonality between tumour and blood are distinct

Using the matched blood and tumour samples, we observed that the clonality of T cells between blood and tumour is highly divergent (**Supplemental Figure 3d**). Within the blood CD4 T cell compartment, only CD4 senescent and Th1 T cells had high levels of intra-subset clonality. We noted that Tregs were not clonal in blood, with <5% of these cells comprising expanded clones, a possible indication that Tregs from TME expanded clones are tissue resident. In the CD8 T cell compartment, the CD8 EMRA and senescent populations were the most clonal populations and are significantly more clonal than their corresponding tumour T cell counterparts (**Supplemental Figure 3e**); which is to be expected for these populations as they are driven through chronic antigen exposure (in many cases viral)^29^. There was no difference in CD8 T cell clonality observed in the blood between ME and AE patients (**Supplemental Figure 3d**). Finally, to determine if the T cell responses within the tumour were enriched for systemic anti-viral responses rather than potential tumour-specific responses, we screened the tumour and blood-derived TCRs against a library of known anti-viral TCRs (see **Methods**). Indeed, we found that anti-viral T cell clones are not enriched in the tumour compared to the blood and there were no widespread consistent differences between ME and AE patients (**Supplemental Figure 3f**). This supports that tumour clones are not enriched for systemic non-tumour-reactive clones.

### Immunosurveilling and resident tumour-infiltrating B and T cell clones are phenotypically distinct

Our dataset benefits from having matched blood and tumour samples taken at the same timepoints allowing us to perform analysis to identify circulating tumour infiltrating lymphocytes (TILs). These can be identified from clones shared between blood and tumour and represent clonally expanded lymphocytes recirculating between tumour and blood and will therefore be critical for immunosurveillance. We identified B and T cells clones that were (a) shared between blood and tumour (recirculating clones), (b) tumour-only (non-circulating TIL clones) and (c) blood-only clones. These states, by definition, exhibit different cellular tendencies for tumour ingress and/or adhesion (**Figure 4a**). Circulating TIL B cells were enriched for naïve B cells in both AE and ME groups, suggesting that naïve B cells may be major components of immunosurveillance and selected B cells become activated in response to the TME (**Figure 4b**). This is supported by the elevated IGHM usage in circulating TIL B cell clones (**Supplemental Figure 4a**). Non-circulating TIL B cells were enriched for antibody secreting B cells and activated memory, suggesting that these are much less mobile upon tumour entry or differentiation. The dynamics of immune cell infiltration is explored in the next section.

**Figure 4:**
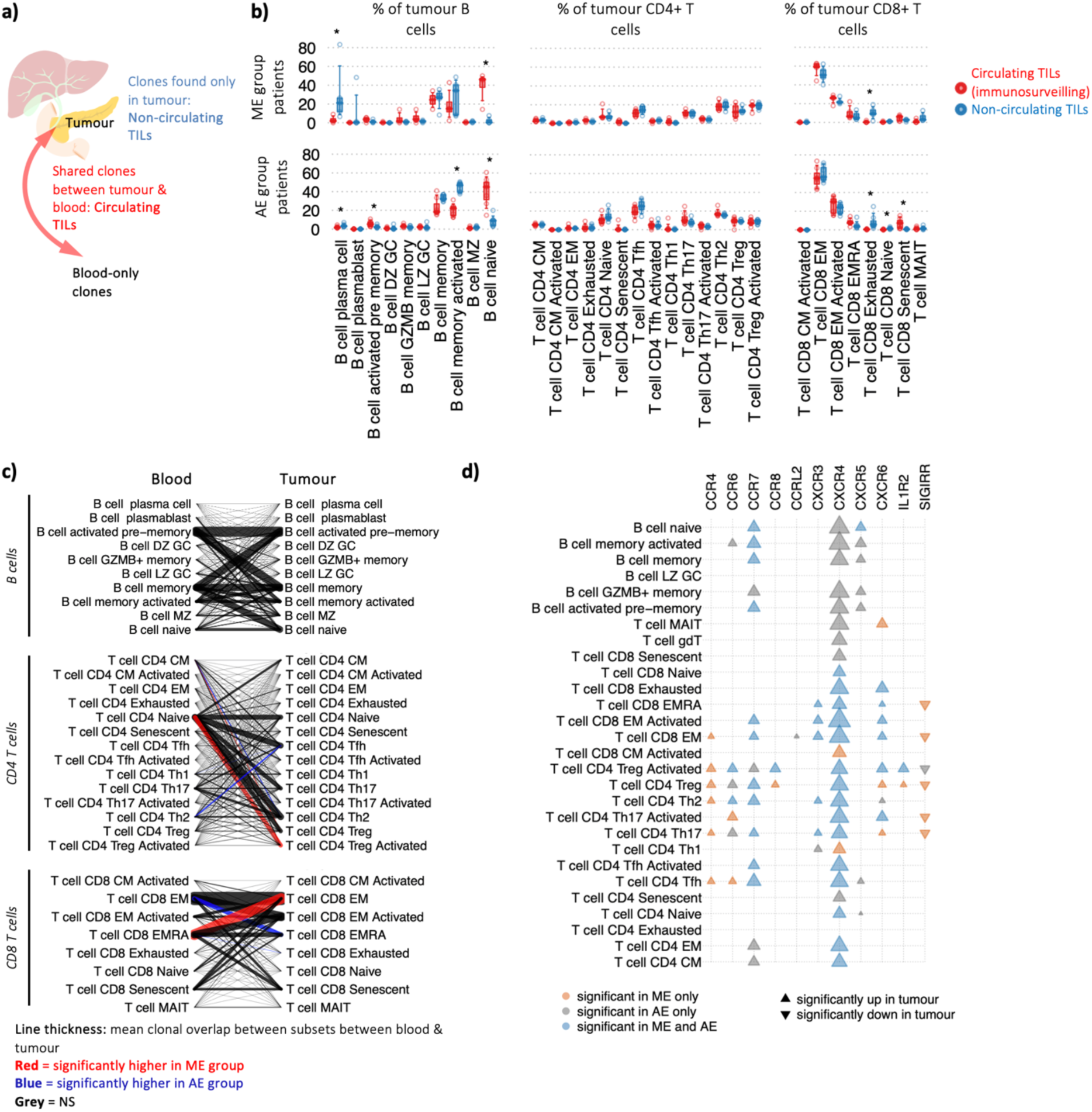
Immunosurveilling and resident B and T cell clones are phenotypically distinct. a) Schematic of clonal definitions: B and T cells clones that are (a) shared between blood and tumour (recirculating clones), (b) tumour-only (non-circulating TIL clones) and (c) blood-only clones. b) The percentage of tumour B cells, CD4 T cells, and CD8 T cells that (red) have clonal members in the blood or (blue) no clonal members in the blood for the ME patients (top) and AE patients (bottom). c) Clonal migration overlap plot, showing the linked phenotypes between blood and tumour B and T cells from shared clones between sites. Line thickness represents the relative means calculated over each patient. Red lines indicate that the corresponding clonal sharing between the corresponding cell types is significantly higher in the ME patients than AE, and a blue line denotes that the clonal sharing between the corresponding cell types is significantly lower in the ME patients than AE. d) Heatmap of DGE between blood and tumour biopsy between ME and AE patients per lymphocyte cell type. For each chemokine receptor and for each cell type, the upwards triangle denotes significant elevation of expression in tumour compared to blood and downwards triangle denotes significant reduction of expression in tumour compared to blood. The triangles are coloured orange, grey and blue if the significance is observed in ME patients only, AE patients only or both, respectively. The sizes of the triangles denotes relative mean expression. All analyses in this figure were performed on the PancrImmune dataset. * denotes p-values<0.05, and tests were performed by MANOVA.

Next, we explored the dynamics of immune cell infiltration. Whilst there was no significant difference of specific CD4 T cell subsets between circulating and non-circulating TILs, circulating CD4 TILs are dominated by Tregs, Tfh, and Th2 (**Figure 4b**). Circulating CD8 TILs are dominated by CD8 EM T cells, which is consistent with the arrival of activated CD8 T cells from the tumour-draining lymph nodes. However, these were not statistically enriched compared to non-circulating CD8 T cell clones only found in the tumour. As expected, exhausted clones were enriched in the TME where they are most likely to encounter their antigen.

We did not observe differences in the proportions of total recirculating B and T cell TIL clones between ME and AE patients (**Supplemental Figure 4b**). However, recirculating B and T cell TIL clones were significantly more expanded than clones private to the tumour or blood clones (**Supplemental Figure 4c**). Finally, through screening the TCRs with a library of known anti-viral TCRs, we find that anti-viral T cell clones are not enriched in the circulating TILs compared to the non-circulating and blood-only T cell clones (**Supplemental Figure 4d**). This supports the conclusion that recirculating TIL clones are not enriched for systemic non-tumour-reactive clones.

### Dynamics of recirculating tumour-infiltrated B and T cells

Next, we considered how the phenotype of clonally-related B and T cells differ between the blood and the tumour. This can be measured through determining the phenotypes of B and T cells within the same clone shared between the blood and tumour (**Figure 4c, Supplemental Figure 4e-g**). Many of the recirculating B and T cells have different phenotypes between blood and tumour, suggesting extensive intra-tumoural B and T cell differentiation within the tumour site and/or distal from the tumour. For B cells, the majority of recirculating B cells are derived from tumour-infiltrated naive, memory and activated pre-memory B cells. This suggests that selected naive B cells from the blood infiltrate the tumour, and these differentiate to express memory B cell markers before recirculating. This also provides evidence that the tumour may be a major site of B cell activation.

For CD4 T cells, the largest overlap occurs between CD4 naïve and T helper phenotypes (Tfh, Th17, Th2, and Tregs) (**Supplemental Figure 4f**), suggesting naïve CD4 cells are being polarised based on intra-tumoural factors. The ME patient group has significantly higher overlap between naïve CD4s and activated Tregs, supporting that the myeloid-enriched TME in these patients is driving the differentiation and proliferation of activated Tregs from naïve cells. Blood CD8 senescent are predominantly related to CD8 EM, activated EM, EMRA and senescent, suggesting that these cells are derived from highly activated effector T cell clones, as expected^30^. Indeed, the clonal relatedness of blood CD8 EMRA and tumour CD8 EM T cells is supported by the observation that these subsets are the most clonal populations in the blood and tumour CD8 populations, respectively (**Supplemental Figure 2h**). Overall, these results demonstrate that the TME can differentially shape the B and T differentiation in the two patient groups.

### B and T cell infiltration is dependent on chemokine receptor upregulation

Previous reports have shown that chemokines are critical for the infiltration and egress of immune cells from tumours, including the CXCR4-CXCL12 axis shown in mouse models of melanoma^31^, as well as a pre-requisite for the formation of TLSs, including the CXCR5-CXCL13 axis^32^. Therefore, we considered the expression of key lymphocyte chemokine receptors upon infiltrating into the tumour which is possible to assess between matched tumour and blood samples in the PancrImmune dataset where this is possible. The chemokine receptors CCR6, CCR7, CXCR3, CXCR4, CXCR5, and CXCR6 have the highest expression across lymphocytes (**Supplemental Figure 5a**), and CCR8, a known hallmark of tumour infiltrating Tregs, is exclusively expressed on Tregs^33^. We observe significant correlations between some chemokine receptors and lymphocyte infiltration, including CCR8 expression correlating with activated Treg levels (**Supplemental Figure 5b**). Differential gene expression (DGE) between blood and tumour infiltrating B and T cell subsets (see **Methods**) revealed that multiple chemokines and their receptors are upregulated upon entry to the tumour (**Figure 4d, Supplemental Figure 5c**). Upregulation of chemokine receptors in TILs implies that they are central to recruitment and maintenance of these immune cell types within the tumour. Of note, CXCR4 was significantly upregulated across 24 out of 28 lymphocyte populations in the AE patient group, but in the ME group, CXCR4 was not upregulated in B cells, MAIT, gamma/delta or CD8/CD4 senescent T cells, in accordance with their lower prevalence in this patient group. Similarly, CXCR5 was only observed to be upregulated in tumour non-naive B cells and Tfh T cells only in the AE but not in ME patients. Lower CXCR5 in tumour B cells and Tfh T cells in ME patients will likely impact the effectiveness of B cell migration, retention and responses within the tumour site. CCR8 expression is significantly increased in intra-tumoral Tregs compared to blood, predominantly in ME patients, with the highest CCR8 expression observed in activated Tregs. Indeed, the same trends were observed when considering only immunosurveilling clones (clones shared between blood and tumour) (**Supplemental Figure 5d-e**). Overall, we observed reduced chemokine receptor expression in intra-tumoural ME patient lymphocytes.

Finally, we show that only TIL T cell clones tend to be acutely activated with elevated with CD69 and PD1) whereas the blood counterparts of same clones are not acutely activated (**Figure 4d**). However, the significant upregulation of CD69 was observed in only AE patients for B cells and multiple CD4 T cell populations, suggesting reduced activation in specific lymphocyte subsets in ME patients.

### Myeloid cells in ME patients dominate cell-cell communication

We have thus far shown that differential immune cell subtype frequencies distinguish ME and AE patients and lymphocyte-associated differences. We next examined the cell-intrinsic differences in cell-cell communication between immune cells between ME and AE patients. Here we considered cell-cell interaction strengths between known cytokine- and inflammation-associated ligands and their receptors (see **Methods**). The signalling strengths between each pair of cell subtypes for each receptor-ligand pair was calculated by multiplying the percentage of cells per cell subtype expressing each respective gene for each patient. Thus, the strengths are independent of the total proportions of each cell type within the tumour (**Figure 5a**). The cell-cell communication network depicted an expected high level of complexity within the tumour microenvironment with each cell subtype providing and receiving signals from many other cells. However, within this complexity, several features were clearly observed. Firstly, ME patients had significantly higher levels of signalling between myeloid and T cell populations, and AE patients had higher levels of signalling between B cell and T cell populations. Indeed, enumerating the number of incoming and outgoing interactions (corresponding to or from cell-surface receptors, respectively, **Figure 5b**) clearly demonstrated that immune signalling within the tumour was dominated by myeloid cells in ME patients and B and T cells in AE patients. The highest levels of incoming and outgoing interactions within ME patients were from MoMac, moDC and granulocyte populations, whereas the highest levels within AE patients were from CD8 EM T cells and memory B cells. Although the proportions of GC and MZ B cells were very low (<5% of total B cells, **Figure 2a**), we observed that these cells have considerable contributions to cell-cell signalling, and indeed significantly higher interactions were seen in the AE patients.

**Figure 5.**
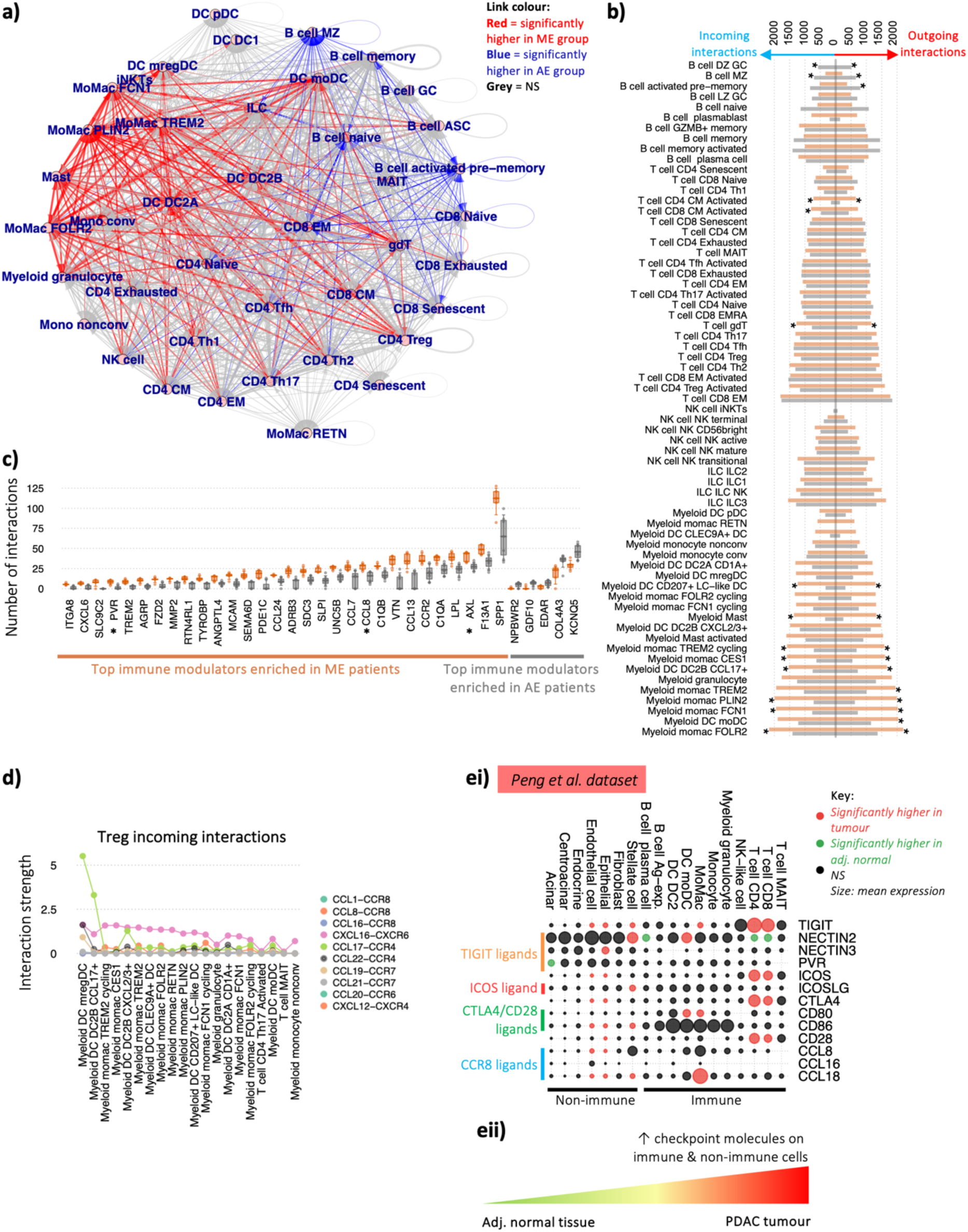
Distinct regulatory mechanisms between patients with different immune cell infiltration. a) Intercellular immune modulator communication network between intra-tumoural immune cells, where each line thickness corresponds to the mean number of receptor-ligand interactions between the corresponding pair of cell types. A red line denotes that the number of receptor ligand-interactions between the corresponding cell types is significantly higher in the ME patients than AE, and a blue line denotes that the number of receptor ligand-interactions between the corresponding cell types is significantly lower in the ME patients than AE. b) Quantification of the number of incoming and outgoing interactions per cell type split by ME and AE patient groups, calculated as a sum of all receptor-ligand pairs identified between all cell types. Bars indicate the means for each patient group, and * denotes p-values<0.05 between groups. c) The number of interactions of the top 30 significantly enriched cytokines in ME patients and all the top 30 significantly enriched cytokines, chemokines and immune-modulators in AE patients (p-values<0.05). d) The top 20 ranked interaction strengths between the key tumour Treg receptors (CCR4, CCR8, CXCR4 and CXCR6) and their ligands per cell type, coloured by receptor-ligand interaction type. ei) Differential checkpoint gene expression between adjacent normal pancreatic tissue and PDAC in both immune and non-immune cell compartments (using the Peng et al. dataset). Red circles indicate significantly higher expression in the tumour and green circles indicate significantly higher expression in the adjacent normal pancreatic tissue. Circle size indicates relative mean gene expression per cell type. ii) Schematic of the checkpoint expression landscape between healthy and pancreatic tumour tissue. All analyses in this figure were performed on the PancrImmune dataset unless otherwise indicated. * denotes p-values<0.05, and tests were performed by MANOVA.

The top significantly enriched immune modulator in the ME patients was SPP1 (**Figure 5c**) which encodes for osteopontin, and is overexpressed in PDAC and known to potentiate tumour cell stemness, M2 macrophage polarisation^34^, checkpoint expression^35^ and is associated with poorer survival across multiple cancers including PDAC^36^. AXL was the third most ME-enriched cytokine that induces mregDC formation and upregulation of PD-L1 expression^37^. Indeed, the top 30 significantly enriched immune modulators in ME patients included CCL8, a ligand for the tumour infiltrating Tregs chemokine receptor CCR8, PVR, a ligand for the T cell checkpoint protein TIGIT and ITGA8, which is known to activate TGFb (**Figure 5c**).

We next considered the signalling interactions to Tregs which are known for being associated with immunosuppression within the tumour microenvironment. The ranked interaction strengths between the key Treg chemokine receptors (CCR4, CCR8, CXCR4 and CXCR6) and their ligands per cell type (**Figure 5d, Supplemental Figure 6a-b**) showed that the incoming interactions with Tregs were dominated by myeloid cells, notably mregDCs (driven by their expression of CCL17 which interacts with CCR4 from Tregs) as well as the regulatory axis CCL22-CCR4 which promotes Treg function^38^, and DC2B CCL17+ and multiple MoMac populations (driven by their expression of CXCL16 which interacts with CXCR6 from Tregs). Finally, interactions with the Treg-exclusive receptor CCR8 were dominated by MoMac expression of CCL8. Indeed, MoMacs were more numerous in ME patients (**Supplemental Figure 1a**), and thus would support the infiltration of Tregs into the tumour region. Whilst the expression of the Treg-associated chemokines, CCL17 and CXCL16, was observed in non-immune cells, including epithelial cells (which includes the tumour cells) (**Supplemental Figure 6c**), the highest expression of these chemokines was within the myeloid MoMac and DC populations. Together, these findings demonstrate a key role of myeloid cells in promoting the immune-regulatory nature of the PDAC TME.

### Checkpoint genes are upregulated across both immune and non-immune cell subsets in PDAC

Finally, we examined the immunosuppressive nature of the whole TME including non-immune cells. Differential gene expression (DGE) analysis between PDAC and normal adjacent tissue from the Peng *et al.* dataset showed significantly elevated checkpoint gene expression in both immune and non-immune cell compartments (**Figure 5ei, Supplemental Figure 6d**). While T cells are the primary source of TIGIT expression, stellate, epithelial and endothelial cells also have increased expression in the tumour compared to pancreatic adjacent normal tissue. Likewise, ICOS and CTLA4 are primarily expressed by T cells, but are significantly higher in expression in tumour compared to pancreatic adjacent normal tissue in epithelial and endothelial cells. Differential expression of TIGIT, ICOS and CTLA4 ligands were observed in both immune and non-immune cell types. Treg-associated chemokine receptor CCR8 ligands, CCL8, CCL16 and CCL18, were also elevated in tumour tissue stellate, epithelial and endothelial cells. Although the highest levels of CCL18 was expressed by MoMacs, stellate cells also contributed significant levels of Treg-specific chemokines suggesting a key role for both immune and non-immune components in shaping the TME into an immunosuppressive environment during tumourigenesis (**Figure 5eii**)^39^.

## Discussion

Our work sheds new light on the potential mechanisms that might underlie the observed differences between myeloid-enriched and adaptive-enriched PDAC tumours. Combining scRNA-seq, CITE-seq and TCR and BCR repertoire analysis of matched blood and tumour samples allowed, for the first time, the identification of different patient groups with distinct immune cell infiltration, selection, differentiation and response mechanisms within the TME, providing a rational way for the selection and design of novel immunotherapeutic interventions for PDAC patients. To this end, we developed several new bioinformatics tools, including (a) *SVMCellTransfer* which allows for efficient and effective annotation of published scRNA-seq datasets based on a reference high-confidence dataset, (b) *scClonetoire* which quantifies the intra- and inter-subset clonality and other repertoire metrics run on single cell multi-omics repertoire data, (c) *scRepTransition* which quantifies the clonal overlap between B or T cell subsets within a sample or between samples. Importantly, *scClonetoire* and *scRepTransition* account for sampling depth differences between samples thus ensuring the ability for statistical comparisons between samples. This comprehensive analysis of the immune landscape within treatment-naive PDAC patients provides a valuable scMulti-omics dataset with high-confidence annotations and important insights into the TME.

Through our multi-omics analyses, we show that dominant immune mechanisms within AE patients, characterised by a low infiltration of myeloid cells and increased proportion of lymphocytes, include dysfunctional GC (or TLS) responses, lower isotype switching, and higher occurrence of IgM+ B cells, and lower generation of plasma cells. The predominance of intra-tumoural memory B cells and the elevated cell-cell interaction signals between B and T cells suggests an antibody-independent role of B cells, such as antigen presentation. Indeed, the predominant contributors of professional APCs in AE patients were B cells, highlighting a potential role for B cells in PDAC TME in shaping T cell activation^40,41^. However, the poor class switching and SHM in AE patients is indicative that some factors that are needed for TLS formation and cell recruitment could be defective, hampering the full development of a GC-like response. T cells, on the other hand, showed clearly increased clonality and higher levels of cytotoxic T cells, yet their ability to control tumour was limited, possibly due to poor infiltration or retention outside the tumour core due to high stromal expression of the CXCR4 ligand, CXCL12 or/and the development of senescence.

ME patients have exhibited a poorer survival signature across multiple studies, and are characterised by the higher infiltration of myeloid cells. Here we showed that ME tumours have a higher presence of immuno-regulatory moMacs and mregDCs. Indeed, we show in our clinical cohort that myeloid cells can act as coordinators of further immuno-suppressive mechanisms through extensive cell-cell signalling mechanisms that are distinct from AE patients, including the attraction of regulatory T cells into the TME^42^. Tregs in ME patients are highly expanded from naive T cells, likely driven by the high levels of TGFb in the pancreatic TME^43,44^. Their association with Tfh T cells could be a driver of differentiation of Tfr (T follicular regulatory cells) which further limit the GC B cell responses in those patients^45,46^. Reduced GC B cell clonal expansion but increased plasma cell fate in ME patients points to direct macrophage-plasma cell cross-talk inducing plasma cell differentiation^22^. GC B cells in ME patients were not as clonal as in AE patients, however, the plasma cell responses are predominantly via IGHA1 and IGHA2. Indeed the IgA isotype can engage with the inhibitory Fc receptor FCAR on myeloid cells, and can mediate inhibitory effects on many immune cell subsets via activation of FcαRI receptors and induction of IL10 production^47^. Indeed, previous studies have shown that tumour-associated antibodies may also exert a pro-tumoural effects through inflammation initiation and maintenance, tissue remodelling, and angiogenesis^48,49^.

We further reveal differential B and T cell selection within the tumour between ME and AE patients. Their infiltration into the tumour may also be limited in ME patients due to the lack of upregulation of key chemokine receptors upon entry into the tissue, including CXCR4 and CXCR5, which have been shown to be important for control of B and T cell trafficking into tissues and play central roles in orchestrating the adaptive cell functions^50^. Indeed, CXCR4 upregulation is known to be driven by factors including hypoxia (HIF1A and VEGF)^51^, where the pancreas is a significantly more hypoxic environment than the blood^52^. However, previous studies have shown that extent of hypoxic areas within the tumour correlates with worse survival of PDAC patients^52^. Alternatively, strong antigen signalling has been shown to downregulate CXCR4 in T cells in melanoma^53^. However, stronger antigen signalling in the ME patients is not well supported due to the higher Treg levels, lower T cell infiltration levels, and lower levels of B-T cell interaction signals. This lays the foundation for future studies determining the key molecular factors influencing lymphocyte infiltration and egress.

Overall, we can identify numerous potential mechanisms that might underlie the observed differences between ME and AE patients and highlight two potential major themes for immune intervention within PDAC patients. In ME patients, targeting the inhibitory myeloid compartment alongside specific targeting of tumour infiltrating Tregs may have the ability to alleviate some of the suppressive mechanisms. For example, the CCR4 and CCR8 pathways are highly active in Tregs and appear to play a crucial role within the TME and their interactions with myeloid cells. It is important to note that the interplay within the TME between the different populations is complex and redundant effects could be at play, as previous reports have suggested that depleting Tregs or fibroblasts can both result in worsening of the disease through the conversion by TGFb of a pathogenic myeloid population even as CD8 T cell responses can be improved^42,54–56^. It is therefore critical to start considering those interventions in combination. AE patient tumours contain diverse lymphocyte subsets, and their activation status suggests that sufficient neoantigens are presented to them. However, the immune-incompetent TME is potentially preventing proper anti-tumour immune responses as can be seen with high levels of CXCR4 on the B cell compartment, potentially restricting their access to the tumour core via retention outside. Indeed, higher number and a specific locations of B cells quality in TME, maturity of TLSs, and neoantigen have been shown in PDAC long-term survivors^16^. These data suggest that patients with higher adaptive cell infiltration may benefit most from boosting the immune response against abortive or dysfunctional TLSs, which may potentially be achieved by cancer vaccines^57^, targeting T cell senescence and/or targeting chemokines. Conversely, patients with higher myeloid cell infiltration may benefit most from selective targeting of Treg functions, such as with anti-CCR8^33,58,59^ and plasma cells.

This study lays the foundation for understanding why immunotherapy has so far not been successful in PDAC and provides an avenue for designing novel therapeutic targets based on a complete understanding of patient intra-tumoural immune heterogeneity. We demonstrate the need for trials to assess changes in immune infiltration over time and under different therapies to build a spatio-temporal understanding of the tumour-immune cross-talk dynamics. Overall, this framework, which combined multimodal data, integrated knowledge-based, unsupervised microstructural annotations, and novel computational tools, has the power to drive niche discovery and can be applied to other tissues in health and disease, such as in cancers with similar AE and ME differential prognostic signatures including glioblastoma^60^, breast^61^, prostate^62^, non-small cell lung cancer, melanoma^63^, bladder cancer^64,65^.

## Materials and Methods

### Sample access and preparation for scRNA-seq

Patients who underwent a curative resection for pancreatic ductal adenocarcinoma were consented for this study. Eight patients were recruited from Oxford under the Oxford Radcliffe biobank (09/H0606/5+5, project: 19/A177). Four patients were recruited from Aachen medical centre under RWTH Aachen biobank project: EK360/19. Informed consent was obtained for all patients. The study was in strict compliance with all institutional ethical regulations. All tumour samples were surgically resected primary pancreatic ductal adenocarcinomas. All tumours were subjected to pathological re-review and histological confirmation by two expert PDAC pathologists before analysis. A supplement providing individual clinical information is provided as Table S1.

The methods for sample collection, PBMC isolation and tissue digestion were previously designed in our manuscript Sivakumar et al. in methods section 5.2-5.4^7^.

### scMulti-omics sequencing and pre-processing

scRNAseq transcriptome processing was performed using the Chromium 10x system involving GEM generation, post GEM-generation clean-up, cDNA amplification and DNA quantification. The library was sequenced using the Illumina NovaSeq platform. Chromium Single Cell Immune Profiling Reagent Kits v1.1 solution was used to deliver a scalable microfluidic platform for digital CITEseq (Cell Surface Protein), GEX, VDJ TCR and VDJ BCR by profiling 500-10,000 individual cells per sample. Libraries were generated and sequenced from the cDNAs and 10x Barcodes were used to associate individual reads back to the individual partitions.

The analysis pipeline applied to process Chromium single-cell data to align reads and generate feature-barcode matrices was performed as previously described^66^. Briefly, gene expression FASTQ files were processed using Cellranger count (v3.1.0) to perform alignment, filtering, barcode counting, and UMI counting, using 10X Genomics’ GRCh38 v3.0.0 reference for Gene Expression analysis and IMGT’s reference for VDJ TCR and BCR analysis. It uses the Chromium cellular barcodes to generate feature-barcode matrices, determine clusters, and perform gene expression analysis.

### Filtering, doublet detection and batch correction of the PancrImmune dataset

For each sample, cells with fewer than 500 transcripts or 500 genes were filtered out. Normalisation and scaling was done using the standard Seurat pipeline. Principal component analysis (PCA) was performed on 5,000 highly variable genes (HVGs) to compute 50 principal components, then *Harmony* was performed (reference) for batch correction, UMAP for dimensionality reduction, and the Louvain algorithm was used for clustering. These clusters were then annotated broadly into B cell, T cell or myeloid clusters based on mapping of >10% BCR+ droplets and elevated CD19 expression, >10% TCR+ droplets and elevated CD3 expression, <10% BCR/TCR+ droplets, respectively.

Doublet identification and removal was performed using both DoubletFinder^67^ and MLtiplet^68^. Each cell type was subsetted into individual objects, and re-clustering within these objects was performed excluding genes which were likely to be influenced by experimental rather than biological factors^69^. These include genes encoding for TCR variable chain, ribosomal proteins, heat shock proteins, mitochondrial proteins, cell cycle proteins, HLA, and noise-related genes (MALAT1, JCHAIN, XIST). For the B and T cell objects, immunoglobulin variable, TCR variable and isotype genes were also excluded.

### Cell type annotations

#### T/NK cell annotations of the PancrImmune data

The re-dimensionality reduced T cell object resulted in 100 clusters generated by k-means. Where ADT-seq data was available this was used in preference to RNA for annotation. T cell clusters were defined by mean proportion TCR expression >0.3, with innate clusters being those with mean proportion TCR <0.3. Individual cells in innate clusters which expressed TRA or TRB sequences were labelled as NK-like T cells.

The innate cells were re-clustered without the T cells to generate 10 clusters, and were labelled by gene expression, ILC1 (TBX21, IFNG, CCL3), ILC3 (RORC, AHR, IL23R IL1R1), gdT (TRDC), NK (EOMES, GZMA, GNLY, KLRC1) based on *de Andrade, et al.*^70^. CD56 bright (immature) NK cells were labelled based on ADT-seq CD56 expression. The remaining NK clusters were labelled based on gene expression patterns to give phenotypic descriptions. NK transitional cells have greater expression of cytokines, chemokines and their receptors (XCL1, XCL2, CXCR4), NK mature cells have greater expression of cytotoxic genes (GZMA, GZMB, PRF1), NK terminal cells have greater expression of adaptive genes (*PRDM1*, ZEB2).

CD4 and CD8 clusters were defined by ADT-seq expression. As has been well documented in T cell single cell papers, there were clusters with overlapping CD4 and CD8 expression. Cells in overlapping clusters were reassigned at the single cell level if either CD4 or CD8 expression was higher. Memory phenotypes were label based on CD45RA, CD45RO, and CD62L expression. Naïve (CD45RA, CD62L), EMRA (CD45RA), EM (CD45RO), CM (CD45RO, CD62L). Further phenotypic labels were based on RNA expression. Exhausted (4 or more of the following: HAVCR2, PDCD1, TOX, LAG3, CTLA4, TIGIT, CD38, ENTPD1). CD4 cells: Treg (FOXP3), senescent (B3GAT1, KLRG1, CD28-, CD27-), Tfh (BCL6, ICOS, CXCR5), Th17 (RORC), Th2 (GATA3), Th1 (TBX21). Finally, clusters were labelled as activated based on HLA-DR ADT-seq expression.

### B cell annotations of the PancrImmune data

The re-dimensionality reduced B cell object resulted in 34 clusters generated by Louvain clustering, and AddModuleScore was used to identify enriched phenotypes (Table S7). Plasma cells were defined as clusters with the percentage of droplets above the 95th percentile BCR nUMIs (percBCR_high) >40% and PC score>0.04, plasmablasts as percBCR_high >15%, naive B cells with >80% unmutated BCRs and >98% IGHD/M BCRs, and memory B cells with mean CD27 expression>0.1. The following cell types were based on AddModuleScores and mean gene expression: B cell memory activated (>0.3 activated score and CD27 expression >0.1), B cell activated pre-memory (>0.4 activated score and CD27 expression <0.1), B cell MZ (>0.8 FGR score and CD27 expression >0.1), B cell GZMB+ memory (GZMB expression>0.3 and CD27 expression >0.1), B cell pre-GC (>0.2 GCB_FT or >0.02 preGC score), B cell GC (>0.3 GC score), of which B cell DZ GC (>0.9 DZ GC), B cell LZ GC (>0.3 LZ GC score). Finally, naive B cells were reassigned at the single cell level if there was >3 SHM, if the isotype was not IGHD/M, or if there was detectable CD27 expression (activated memory) or without CD27 expression (activated pre-memory).

### Myeloid cell annotations of the PancrImmune data

The re-dimensionality reduced myeloid subsetted object was used to identify enriched phenotypes (Table S7). We downsampled the cells to 2000 UMIs/cells and selected variable genes similarly to the seeding step of the clustering. To focus on biologically relevant gene-to-gene correlation, we calculated a Pearson correlation matrix between genes for each sample. For that purpose expression values were log transformed Log(1+UMI(gene,cell) while genes with less than 5 UMIs were excluded. Correlation matrices were averaged following z-transformation. The averaged z matrix was then transformed back to correlation coefficients. We grouped the genes into gene “modules” by complete linkage hierarchical clustering. Specifically, semi-supervised module analysis by complete linkage hierarchical clustering was carried out on variable, biologically-meaningful, and abundantly expressed genes^71^. For example, curated cell-cycle genes and other lateral programs (such as HLA- and HIST-) were excluded from module analysis. Subsequently, myeloid cells were assigned annotations at two levels of granularity based on prior knowledge of marker genes and modules, spanning PDAC and other cancer datasets.

### Annotation of published datasets using SVMCellTransfer

The raw gene-count matrices from Steele et al. and Peng et al. were downloaded from ^10,12^ and filtered using the same parameters as above, and merged. The B, T and myeloid cells were identified and separated in individual objects, merged and batch corrected with the PancrImmune populations via Harmony, and annotated using the custom-written support vector machine (SVM) cell label transfer method, *SVMCellTransfer*. The non-immune cells from the Peng et al. and Steele et al. datasets were merged, batch-corrected and broad cellular annotations were performed using published cell-type markers (**Supplemental Item 1**).

### BCR-seq/TCR-seq analysis

A pipeline, *scIsoTyper*, was written to assign most probable BCR IGH and IGK/L chains per droplet (based on nUMIs) and most probable TCR TRA and TRB chains per droplet (based on nUMIs). Annotations were performed using IMGT, and clonality was performed using a single-cell extension of established VDJ network construction software from Bashford-Rogers *et al.*^24^ as part of *scIsoTyper*.

*scClonetoire* was written to quantify the intra- and inter-subset clonality and other repertoire metrics run on single cell multi-omics repertoire data. Intra-subset clonality measures the number of B, CD4 or CD8 T cell clones with 2 or more cells within each cell subset. This accounts for sampling depth differences between samples through generating a mean across 1000 subsamples a set depth of each sample (n=5 cells). Inter-subset clonality measures the percentage of B, CD4 or CD8 T cells of each cell type as members of clones 3 or more cells across all populations. This accounts for sampling depth differences between samples through generating a mean across 1000 subsamples a set depth of each sample (n=50 cells). These sampling depths were chosen to ensure values were captured across as many immune cell subsets as possible, even when the cell type was rare, whilst still ensuring representation across the sample.

The quantification of clonal overlap between B, CD4 or CD8 T cell subsets within a sample or between samples was performed using a novel pipeline called *scRepTransition*. For the clonal overlap between B, CD4 or CD8 T cell subsets, the absolute number of B or T cells within the same with different cellular annotations was quantified. For all samples in which 1 or more clonal overlaps between cellular annotations was observed, these were normalised to sum to 1. The relative proportions were statistically compared between patient groups via MANOVA.

Viral TCR detection was performed using VDJdb^72^, McPAS^73^ and TCRdb^74^ as reference datasets.

### Cell-cell interaction analysis

The human ligand-receptor database was accessed from Fantom (https://fantom.gsc.riken.jp/), and intercepted the genes that were captured in the PancrImmune scMulti-omics dataset (**Table S5**). The signalling strengths between each pair of cell subtypes for each receptor-ligand pair was calculated by multiplying the percentage of cells per cell subtype expressing each respective gene for each patient. This was calculated for each cell subtype with >=3 cells. This was plotted using *igraph* in R, and MANOVA was used to determine statistical differences between patient groups. The number of inbound and outbound links between cell subtypes was the counts of all corresponding non-zero receptor-ligand signalling strengths (**Figure 5b**). This was computed using *igraph* in R. Ranked interaction strengths per cell type were extracted per receptor-ligand pair for the Treg-specific analysis (**Figure 5c**).

### APC analysis

The pAPC score for each cell (which quantifies the feature expression programme for MHC II and accessory pathway molecules) was calculated using the AddModuleScore using the pAPC pathway genes (Table S7). The distributions of scores from DCs (known pAPCs) and CD8 T cells (known non-pAPCs) was used to define a threshold above which we defined cells as being pAPCs using a logistic regression classifier via fitting a *glm* in R. The statistics between the proportions of tumour pAPC cells between patient groups was performed using MANOVA.

### Differential gene expression analysis and pathway analysis

Pseudobulk differential gene expression methods were employed using the edgeR^75^ package for analysis of aggregated read counts per cell type per patient. This was chosen to reduce the false positive detection rates, reduce biases between patient samples, and address the problem of zero inflated scRNAseq expression data. Briefly, for cells of a given type, we first aggregated reads across cells within each patient. The likelihood ratio test as well as the quasi-likelihood F-test approach (edgeR-QLF). For limma, we compared two modes: limma-trend, which incorporates the mean-variance trend into the empirical Bayes procedure at the gene level, and voom (limma-voom), which incorporates the mean-variance trend by assigning a weight to each individual observation^76^. Log-transformed counts per million values computed by edgeR were provided as input to limma-trend. Differentially expressed genes were defined as adjusted p-values <0.05. Pathway scores per cell were calculated using the *AddModuleScores* function in Seurat in R using pathway gene sets (GO_T_cell_proliferation and REACTOME_apoptosis).

### Survival analysis

Data from the PAAD TCGA (https://portal.gdc.cancer.gov/) was downloaded and normalised. Patients that were not pathologically PDAC including samples with <1% neoplastic cellularity, neuroendocrine, IPMN and acinar cell carcinoma were excluded based on sample annotations (http://api.gdc.cancer.gov/data/1a7d7be8-675d-4e60-a105-19d4121bdebf). R packages survival and survminer were employed. The Kaplan Meier (KM) curve was plotted using *survfit* in R to observe survival probabilities over time between patient groups (high versus low IGHM expression). Surv_cutpoint() and surv_categorize() was used to determine an optimal cutpoint using maximally selected rank statistics for IGHM expression. The cox regression model was used to estimate and compare hazard ratios between IGHM high and low groups.

### Cell composition deconvolution

Deconvolution between the PancrImmune and TCGA datasets was carried out using BayesPrism^77^, a Bayesian method to infer cell type fraction. The intersection of genes present in both datasets was identified, and the raw untransformed count data was used. To assign cell type labels, cell types from annotated single-cell data was used. Substantial heterogeneity was accounted for by creating cell state labels through sub-clustering of cell types within each patient. A threshold of 50 cells per cell state was applied to ensure a sufficient number of cells for accurate sub-clustering. Genes related to ribosomes, mitochondria, chromosome X, and chromosome Y were filtered out from the analysis, as their presence could introduce bias. When running prism object, count matrix was used as input type and key was set to NULL since there were no malignant cells present in the PancrImmune dataset, as recommended by the authors. All other parameters were left at their default values. The mean cell expression was then obtained from get.theta() function. Subsequently, downstream analysis included PCA to divide TCGA cohort as myeloid high, adaptive enriched followed by plotting the proportions.

### Code and data availability

All code is available via https://github.com/rbr1/scIsoTyper/, https://github.com/rbr1/PancrImmune_PDAC_paper, and https://github.com/sakinaamin/BayesPrism. Data will be made available via XX (currently in progress).

## Supporting information

Supplemental item

## Acknowledgements

Firstly, we would like to thank the patients and clinicians who contributed to this study. S.S. was funded on an Oxford-BMS Fellowship with additional funding for the single cells sequencing from BMS and the CRUK Oxford Centre. M.L.D. and J.W. were supported by the Kennedy Trust for Rheumatology Research Cell Dynamics Platform 20 21 17. R.J.M.B.-R. and F.A.T. were supported by the Wellcome Trust, University of Oxford and Oxford Cancer Centre. B.S was supported by the Association of British Neurologists via the Patrick Berthoud Charitable Trust. M.R.M and B.F are supported by the NIHR Biomedical Research Centre at Oxford. S.H. was supported by the NIH NCI transition fellowship K00 CA223043. The views expressed in this article are those of the authors and not necessarily those of the National Health Service, the NIHR, or the Department of Health. A.J. was funded by a Clarendon scholarship and S.A. was funded by Clarendon in partnership with St John’s College. We thank Theodosios Kyriakou and Angela Lee for overseeing the sample submission at the Wellcome Centre for Human Genetics in Oxford. This work was supported by BMS.

## Authors’ contributions

S.S. and E.A-S., M.L.D., R.J.M.B-R conceived and designed the analysis. S.S., L.H., F.T., H.S., S.R., I.N., J.W., A.E.F., G.W., U.P.N., P.C., L.S., R.J.M.B-R collected the data. S.S., A.J., E.A-B, M.N.L., P.K.S., S.A., S.H., A.M., B.S., S.R., I.N., B.F., R.J.M.B-R contributed data or analysis tools. A.J., E.A-B, M.N.L., P.K.S., S.A., S.H., A.M., S.R., I.N., D.P., M.Merad, M.L.D., R.J.M.B-R performed the analyses. All authors contributed intellectual input/interpretation. S.S., A.J., E.A-B, M.L.D., E.A-S., R.J.M.B-R wrote the paper with input from all other authors.

## Ethics approval and consent to participate

Informed consent was obtained for all patients. The study was in strict compliance with all institutional ethical regulations.

---

**Supplemental Figure 1.**
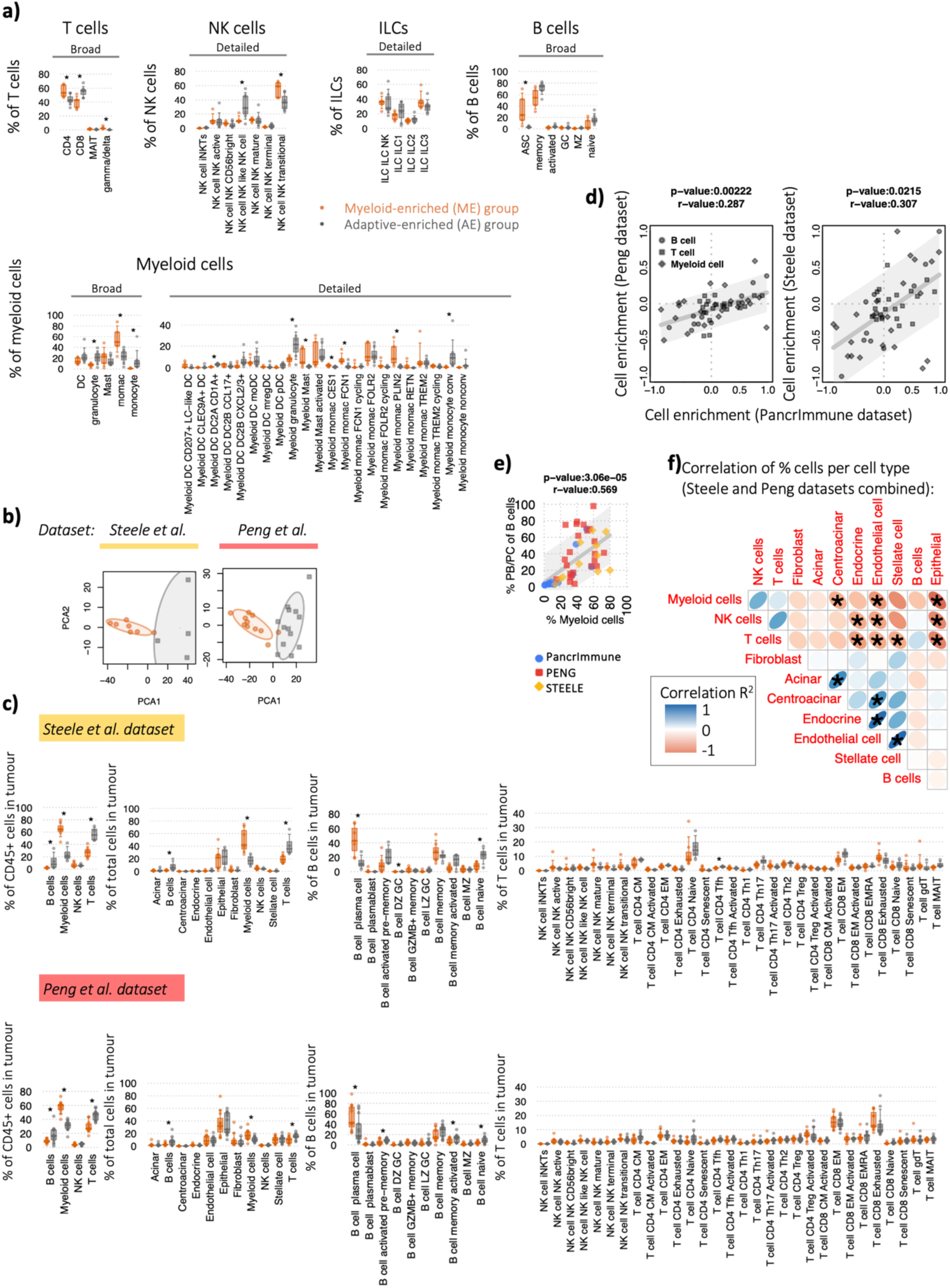
a) Tumour immune cell subset proportions between ME and AE patient groups within cellular subsets as a proportion within the PancrImmune dataset. Orange represents ME patients and grey represents AE patients. b) Principal component analysis (PCA) based on PDAC CD45+ immune cell infiltration proportions for the Steele and Peng datasets, coloured by patient group. c) Cell subset proportions between ME and AE patient groups within cellular subsets as a proportion for the Steele and Peng datasets. Orange represents ME patients and grey represents AE patients. d) Correlation of the cell enrichment between ME and AE patients between the PancrImmune tumour and (left) Peng and (right) Steele datasets. P-values and R^2^ values provided above each plot. e) The correlation of myeloid cells as a proportion of total intra-tumoural immune cells with plasmablasts and plasma cells as a proportion of total B cells, coloured blue, red and yellow for the PancrImmune, Peng and Steele datasets respectively. f) Correlation of immune and non-immune cell proportions from the Steele and Peng datasets combined as a proportion of total cells. The blue positively sloped ellipses represent positive correlations and negatively sloped ellipses represent negative correlations, and * denotes significant correlations p-values<0.05. In panels (a) and (c), * denotes p-values<0.05, and tests were performed by MANOVA.

**Supplemental Figure 2.**
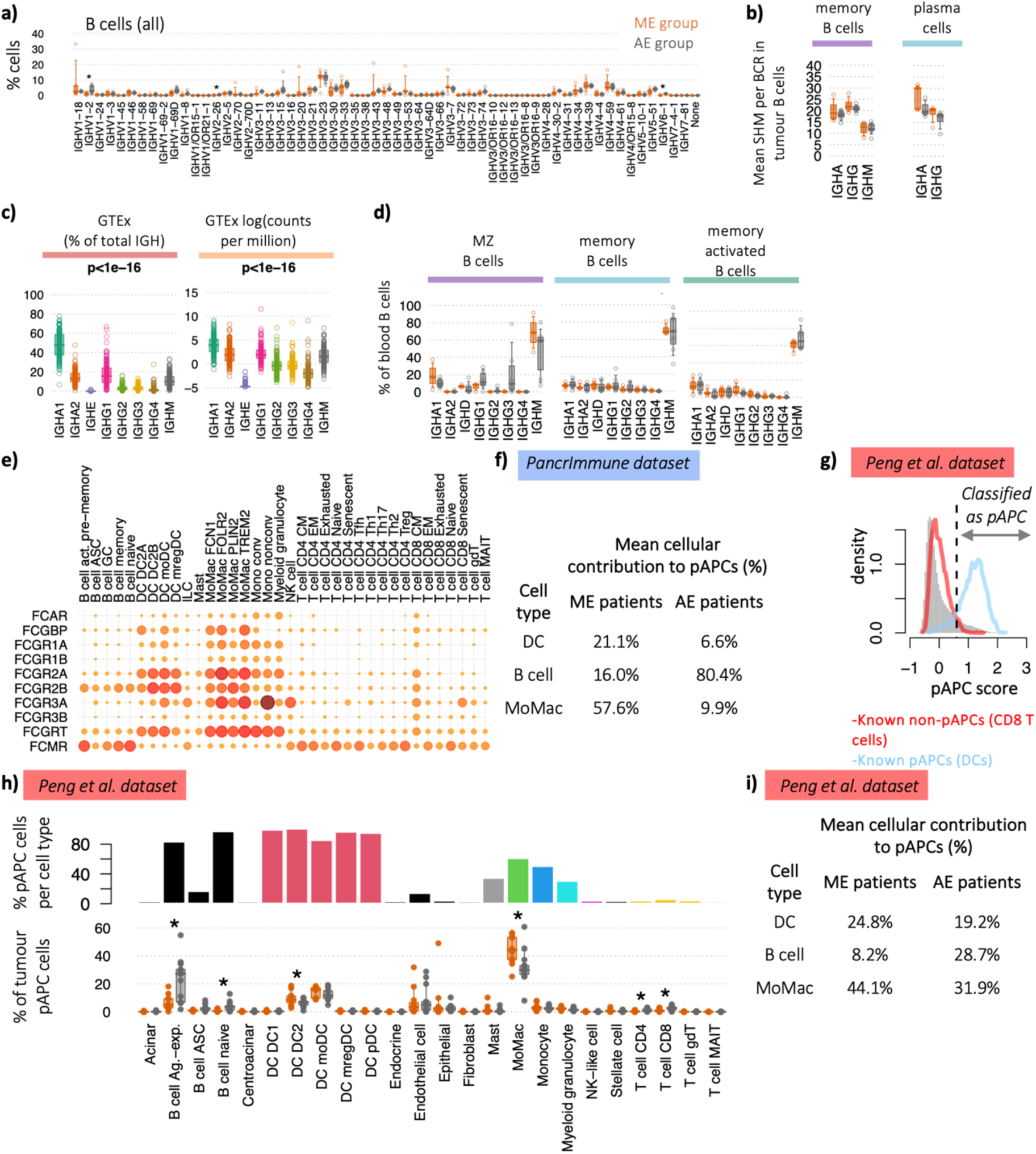
a) IGHV proportions between ME and AE patient groups of total tumour B cells within the PancrImmune dataset. Orange represents ME patients and grey represents AE patients. b) Mean SHM levels between ME and AE patient groups for tumour memory B cells and plasma cells within the PancrImmune dataset. c) Isotype usages (left) as a proportion of total IGH and (right) counts per million in healthy pancreatic tissue from the GTEx RNA-seq dataset. P-values generated by ANOVA. d) The proportions of blood B cells within activated, memory and plasma cells expressing each isotype, coloured by patient group using the PancrImmune dataset. e) The relative levels of the FC receptor gene expression between intra-tumoural cell types, where larger circle size indicates higher expression using the PancrImmune dataset. f) Table of the mean cellular contribution to pAPCs between ME and AE patient groups, using the PancrImmune dataset. g) Histogram of the professional antigen presentation (pAPC) scores for (grey) all cells, (red) CD8 T cells and (blue) DCs, using the Peng dataset. Dashed line indicates the threshold for classification of pAPCs. h) (top) Barchart of the percentages of pAPCs comprising each cell type, and (bottom) the proportion of pAPCs comprising each cell type between patient groups, using the Peng dataset. i) Table of the mean cellular contribution to pAPCs between ME and AE patient groups, using the Peng dataset. * denotes p-values<0.05, and tests were performed by MANOVA.

**Supplemental Figure 3.**
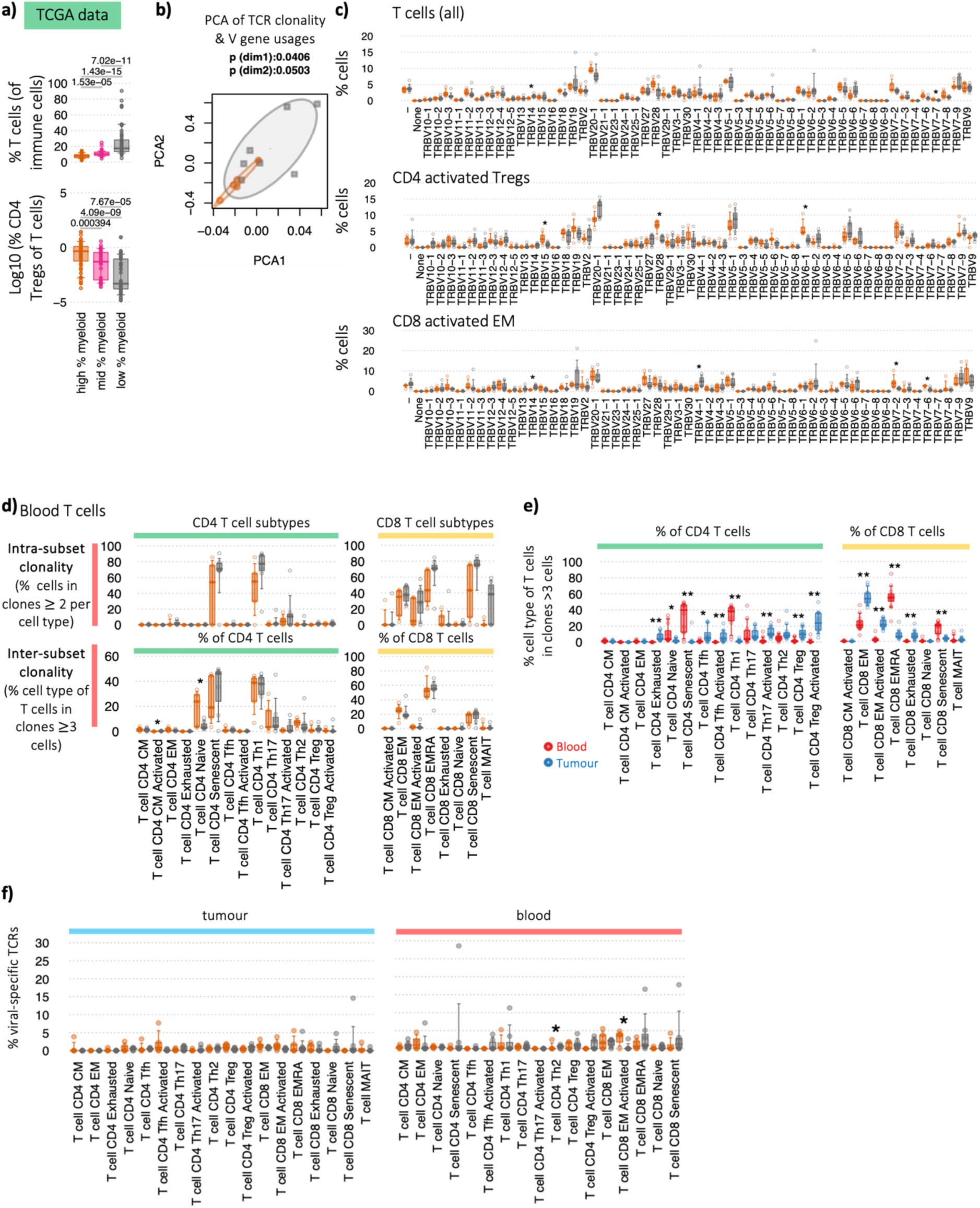
a) Correlation of the Treg proportions (as a proportion of total T cells) with myeloid cell proportions (as a proportion of total immune cells). Cellular deconvolution of the PAAD TCGA dataset (n=156 patients) by BayesPrism using the PancrImmune dataset as a reference. The TCGA patients were split into tertials based on myeloid cell proportions (low % myeloid cells = lowest 33% of patients, mid % myeloid cells = mid 33% of patients, high % myeloid cells = highest 33% of patients). P-values calculated by Wilcoxon test. b) Principal component analysis (PCA) of the TCR clonality and TRB V gene usages, coloured by patient group. c) TRBV proportions between ME and AE patient groups of total tumour T cells, activated Tregs and CD8 activated EM T cells within the PancrImmune dataset. Orange represents ME patients and grey represents AE patients. d) Clonality of the blood T cell subpopulations between the ME and AE patient groups via two measures: (top) *intra-subset clonality* (the percentage of cells in clones >2 cells per subset, measuring the clonality within the subset thus reflecting specific cell populations which are actively expanding), and (bottom) *inter-subset clonality* (the percentage of cells of each cell type as members of clones >3 cells across all populations, demonstrating, this indicates cells within each T cell subset that may be members of larger clones than span multiple phenotypes, reflecting T cell plasticity of expanding clones). e) The *inter-subset clonality* between tumour (blue) and blood (red) T cells. f) The percentage of TCRs from each T cell subset that match to anti-viral T cell clones, coloured by patient group. All analyses in this figure were performed on the PancrImmune dataset. * denotes p-values<0.05, and tests were performed by MANOVA.

**Supplemental Figure 4.**
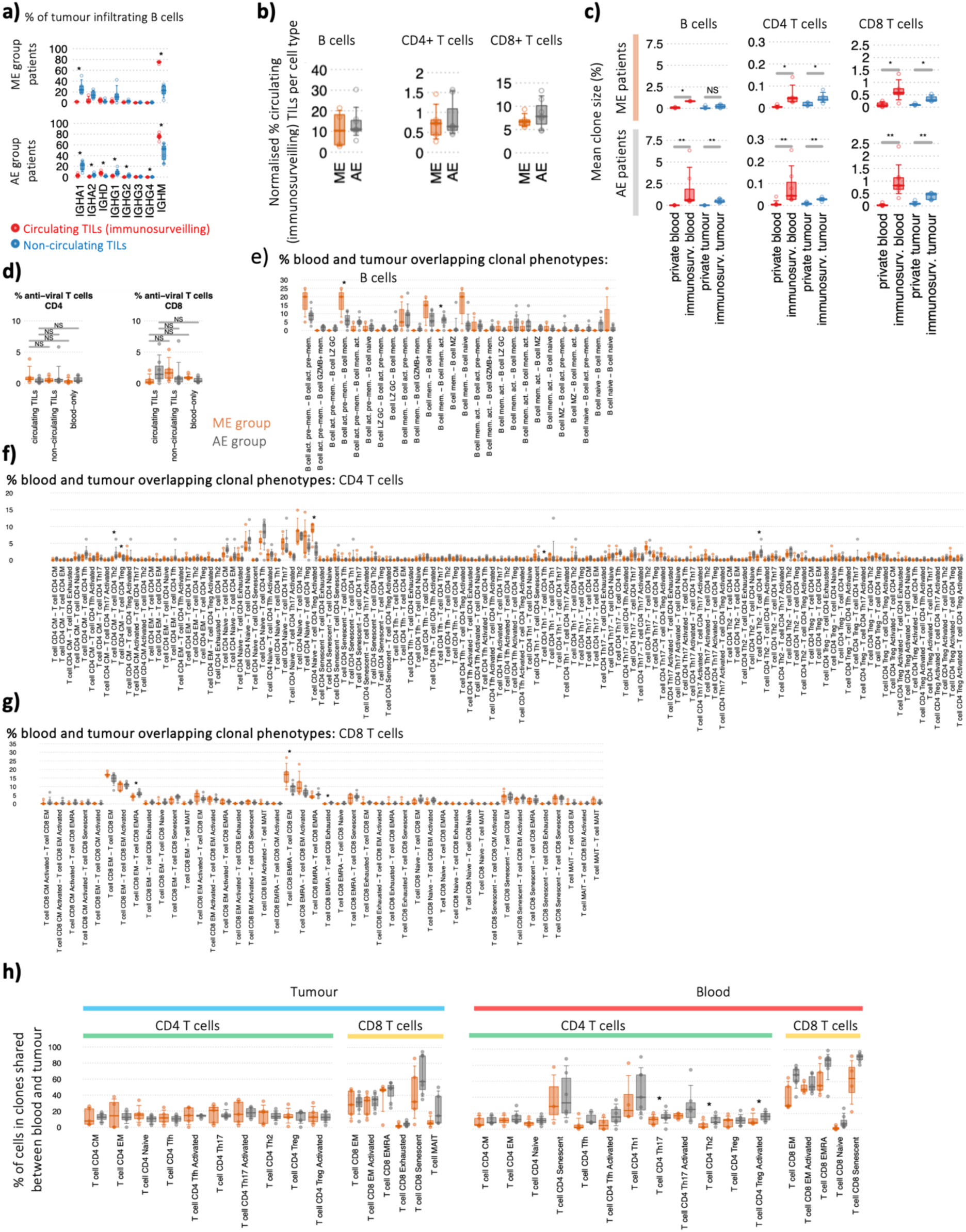
a) The isotype usage percentages of tumour B cells have clonal members in the blood or (blue) no clonal members in the blood between ME patients (top) and AE patients (bottom). b) The normalised level of re-circulating tumour clones between ME and AE patients B and T cell clones. c) The mean clone sizes per patient between blood and tumour re-circulating and private clones, plotted between ME and AE patient groups. * denotes p-values<0.05, ** p-values<0.005. d) The percentage of TCRs that match to anti-viral T cell clones between re-circulating and private clones, coloured by patient group. The relative proportions of clones overlapping blood and tumour split by phenotype for e) B cells, f) CD4 T cells, and g) CD8 T cells, coloured by patient group h) The percentage of cells in clones shared between blood and tumour, split by source and coloured by patient group. All analyses in this figure were performed on the PancrImmune dataset. Unless otherwise mentioned, * denotes p-values<0.05 and tests were performed by MANOVA.

**Supplemental Figure 5.**
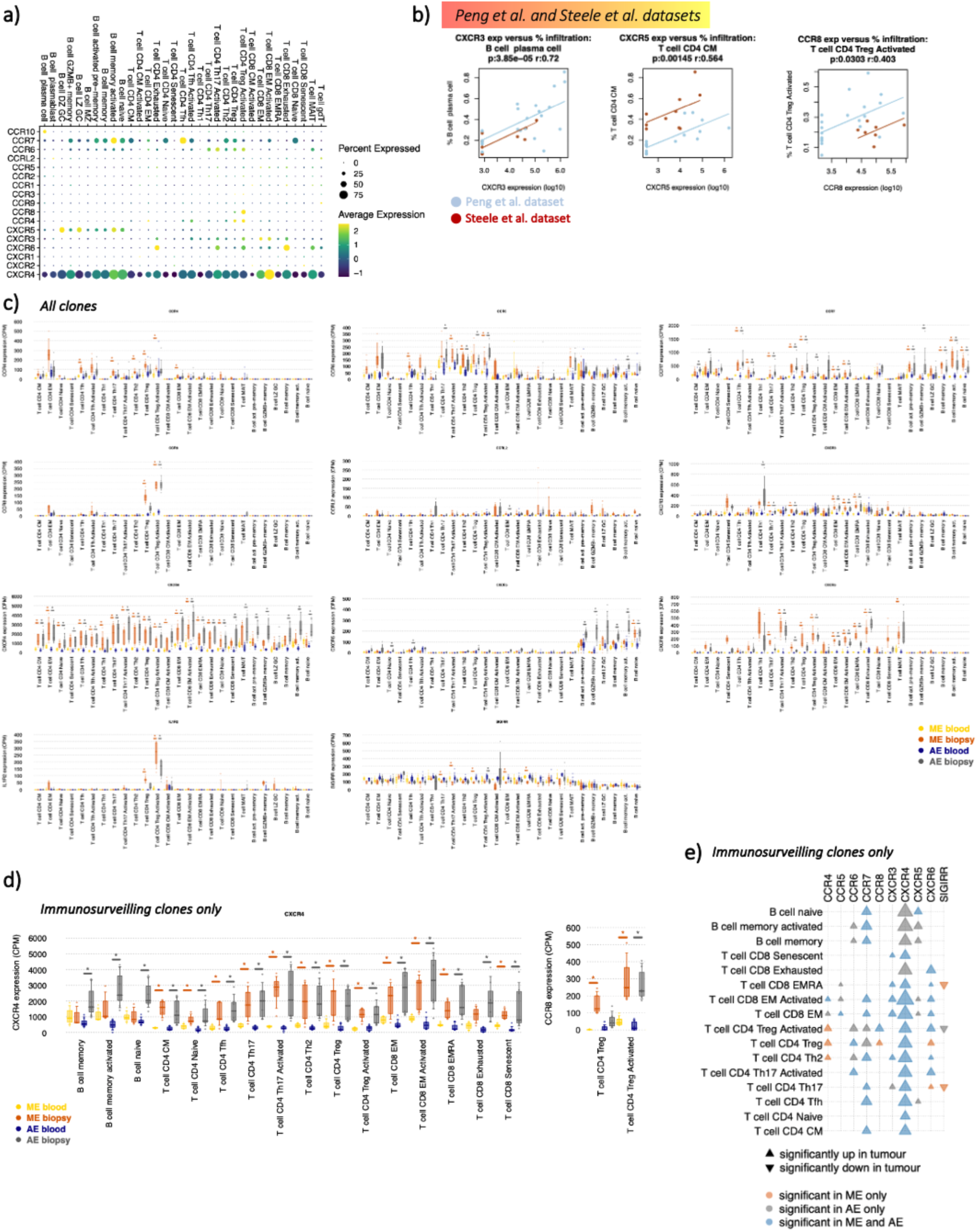
a) Gene expression dot plot of the chemokine receptor gene expression across lymphocytes. b) Correlations between immune cell proportion within tumour microenvironment and mean chemokine expression of that cell type. P-values and r^2^ values were computed using the repeated measures correlation from rmcorr package in R, with the patients from the Peng et al. data in blue and the Steele et al. data in red. c) The mean gene expression of key lymphocyte chemokine receptors between all blood and tumour biopsy between ME and AE patients for each cell type. Each point represents the mean expression per patient per cell group. d) The mean gene expression of key CXCR4 and CCR8 between blood and tumour biopsy between ME and AE patients for each cell type for only immunosurveilling clones (clones shared between blood and tumour). Each point represents the mean expression per patient per cell group. * denotes p-values <0.05 as determined by DGE. e) Heatmap of DGE between blood and tumour biopsy for immunosurveilling clones (clones shared between blood and tumour) between ME and AE patients. For each chemokine receptor and for each cell type, the upwards triangle denotes significant elevation of expression in tumour compared to blood and downwards triangle denotes significant reduction of expression in tumour compared to blood. The triangles are coloured orange, grey and blue if the significance is observed in ME patients only, AE patients only or both, respectively. The sizes of the triangles denote relative mean expression. All analyses in this figure were performed on the PancrImmune dataset. Unless otherwise indicated, * denotes p-values<0.05 and tests were performed by MANOVA.

**Supplemental Figure 6.**
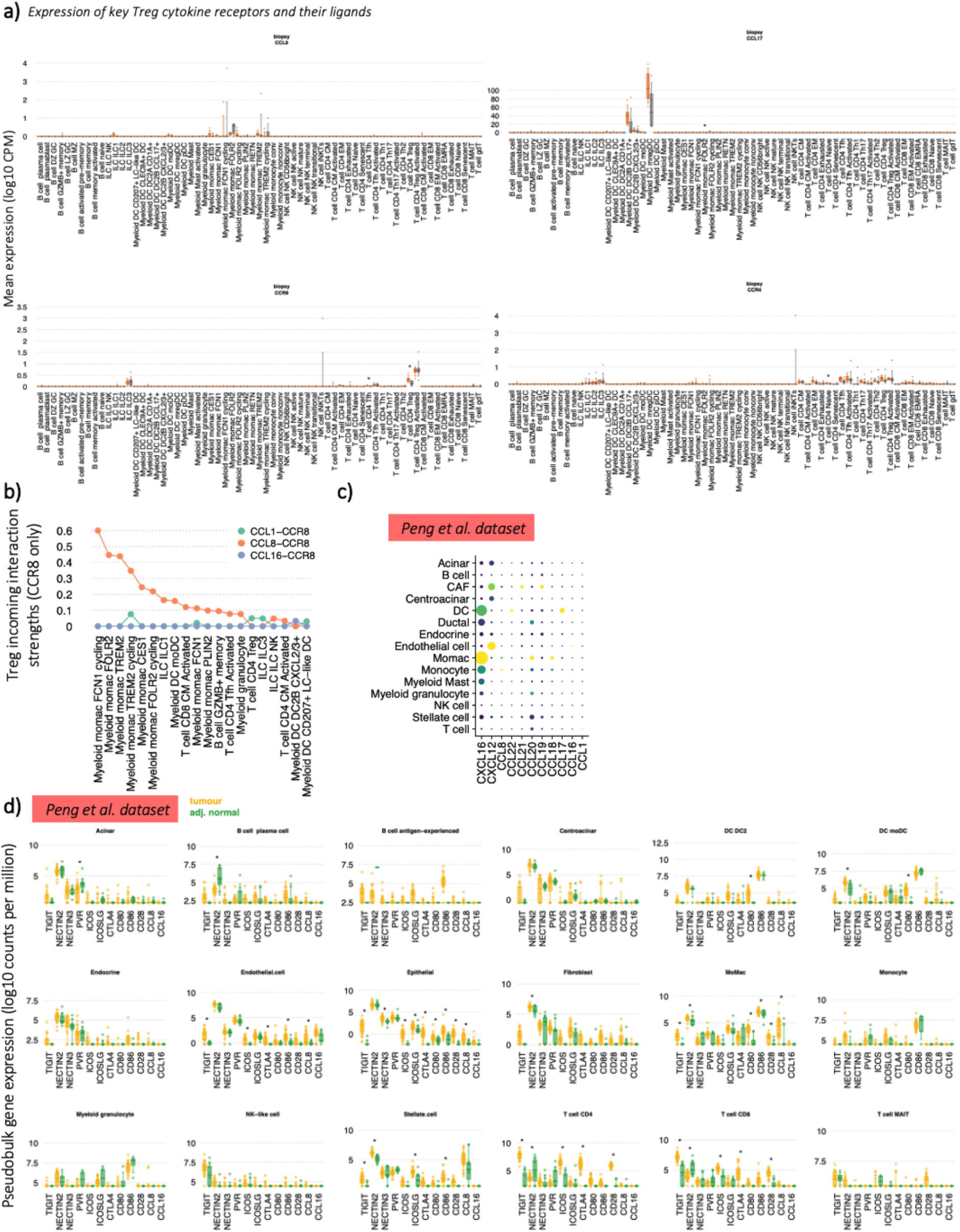
a) The mean normalised gene expression per sample of key Treg chemokines and their receptors across cell types within the tumour from the PancrImmune dataset. * denotes p-values<0.05 and tests were performed by MANOVA. b) The top 20 ranked interaction strengths between the exclusive Treg receptor CCR8 and its ligands per cell type, coloured by receptor-ligand interaction type. c) The relative expression of key lymphocyte chemokines across both immune and non-immune cells, using the Peng dataset. The circle colour denotes relative mean expression level (yellow indicates higher levels) and size indicates percentage of cells expressing each gene. d) Boxplots differential checkpoint gene expression between adjacent normal pancreatic tissue and PDAC in both immune and non-immune cell compartments using the Peng dataset.

