## Supplemental item for "Single-cell immune multi-omics and repertoire analyses in pancreatic ductal adenocarcinoma reveal differential immunosuppressive mechanisms within different tumour microenvironments"

Sivakumar et al.

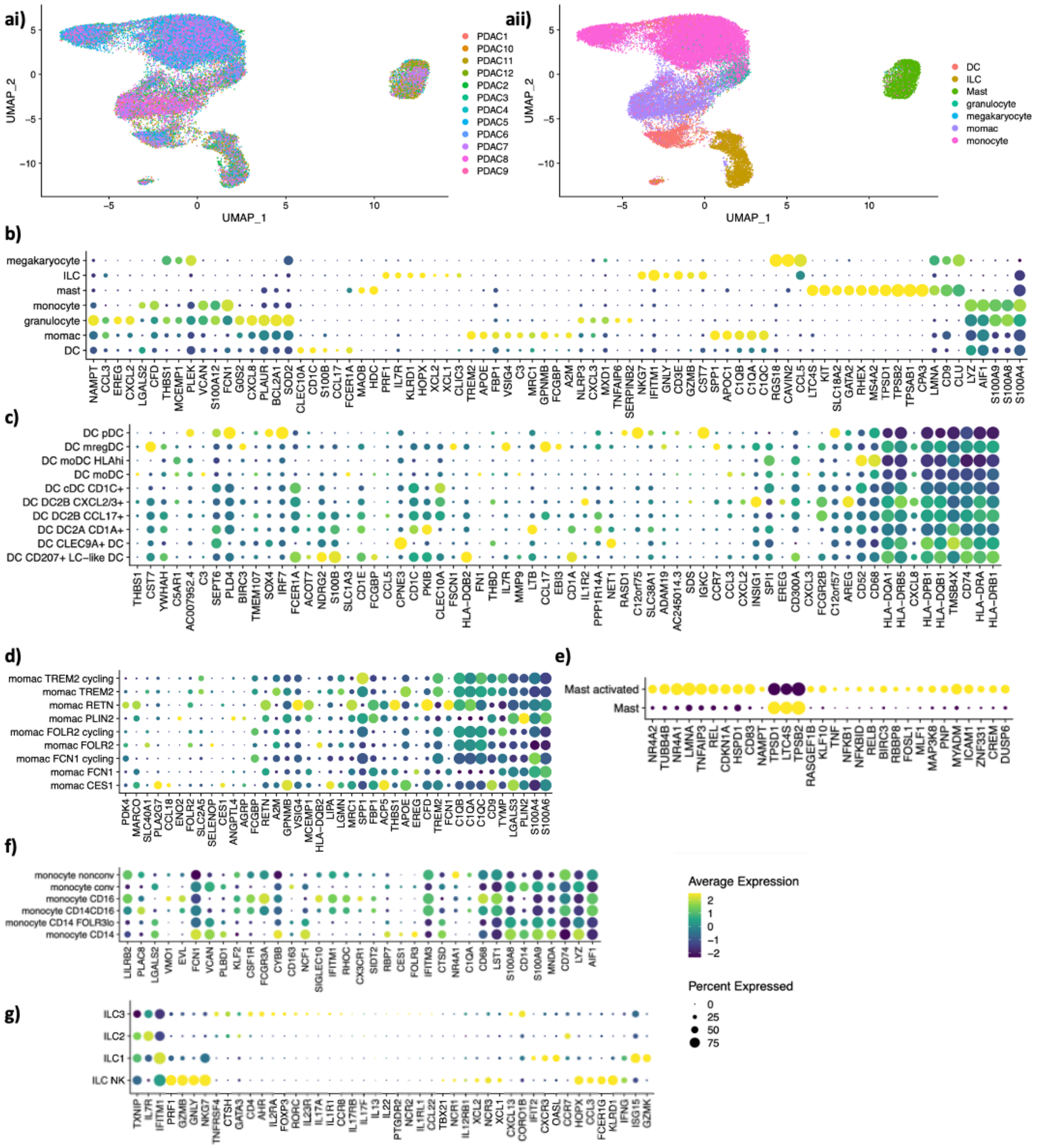

**Supplemental item figure 1.1.** Gene signatures for the myeloid cell annotations from the PancrImmune data.

**a)** UMAP projection of the myeloid cells coloured by (i) patient ID and (ii) broad cellular label.

**b-g)** Gene signatures for the (b) broad annotations, (c) DCs, (d) momacs, (e) mast cells, (f) monocytes, (g) ILCs from the PancrImmune data.

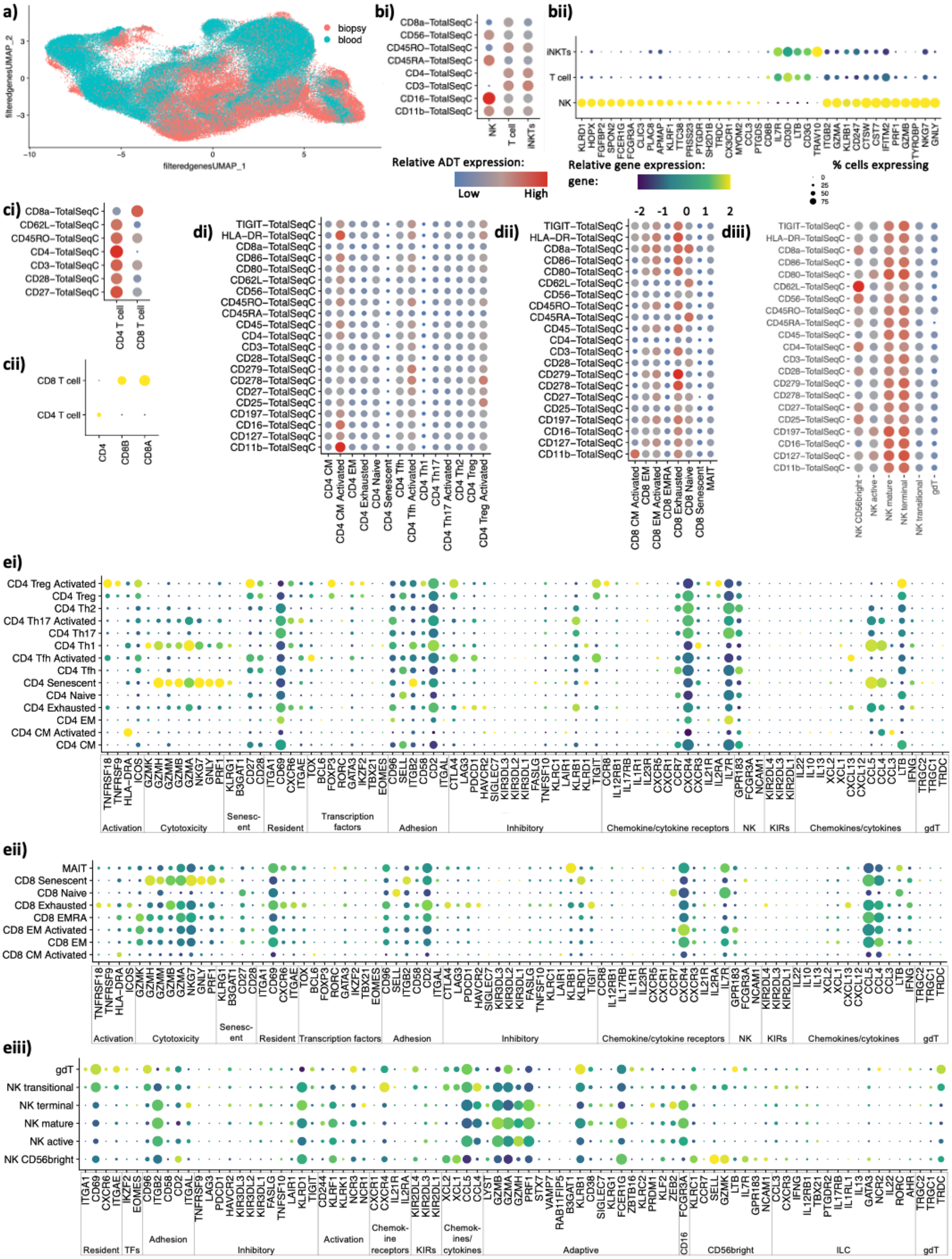


**Supplemental item figure 1.2.** Gene signatures for the T and NK cell annotations from the PancrImmune data.

**a)** UMAP projection of the T and NK cell coloured by sample source.

**b)** Phenotypic signatures for the T and NK cell annotations by (i) CITE-seq and (ii) gene expression.

**c)** Phenotypic signatures for the broad T cell annotations by (i) CITE-seq and (ii) gene expression.

**d)** CITE-seq signatures for the detailed (i) CD4 T cell, (ii) CD8 T cell and (iii) NK cell annotations.

**e)** Gene signatures for the detailed (i) CD4 T cell, (ii) CD8 T cell and (iii) NK cell annotations, grouped by broad gene functional groups.

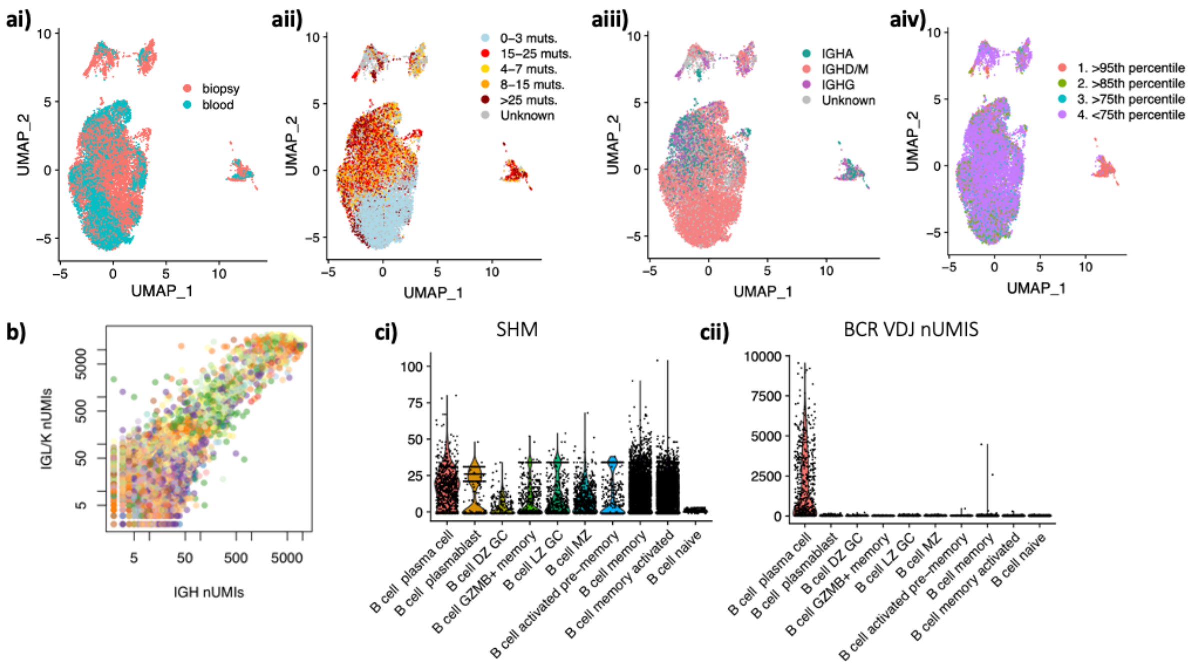


d)

% B cells per isotype per cell type

| | IGHA1 | IGHA2 | IGHD/M | IGHG1 | IGHG2 | IGHG3 | IGHG4 |
|---|---|---|---|---|---|---|---|
| B cell plasma cell | 27.09 | 15.98 | 20.09 | 25.57 | 8.07 | 2.74 | 0.46 |
| B cell plasmablast | 2.21 | 23.53 | 10.29 | 62.13 | 1.10 | 0.74 | 0.00 |
| B cell DZ GC | 2.31 | 0.00 | 42.31 | 43.08 | 3.85 | 7.69 | 0.77 |
| B cell GZMB+ memory | 10.54 | 3.43 | 78.19 | 6.37 | 0.00 | 1.47 | 0.00 |
| B cell LZ GC | 2.42 | 0.66 | 55.16 | 31.87 | 5.71 | 3.08 | 1.10 |
| B cell MZ | 9.58 | 0.63 | 76.45 | 8.01 | 0.63 | 4.71 | 0.00 |
| B cell activated pre-memory | 6.96 | 0.56 | 88.58 | 3.06 | 0.28 | 0.56 | 0.00 |
| B cell memory | 15.76 | 4.25 | 67.18 | 7.90 | 3.41 | 1.10 | 0.40 |
| B cell memory activated | 24.66 | 4.83 | 61.24 | 6.81 | 1.78 | 0.63 | 0.06 |
| B cell naive | 0.00 | 0.00 | 100.00 | 0.00 | 0.00 | 0.00 | 0.00 |

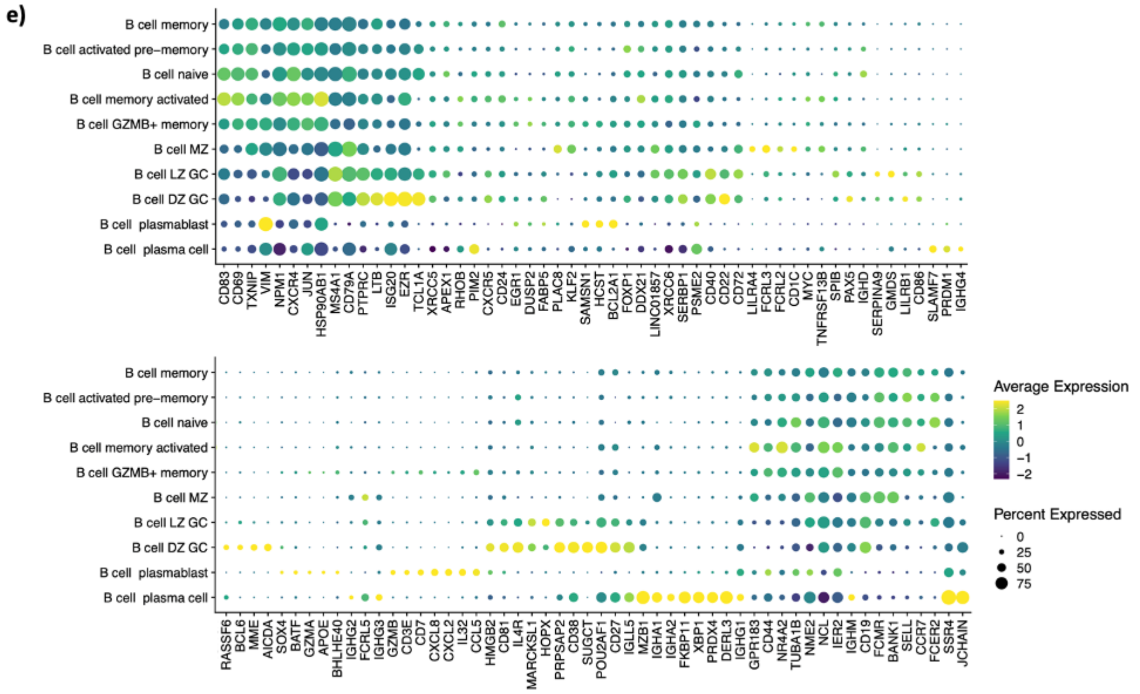


**Supplemental item figure 1.3.** Gene signatures for the B cell annotations from the PancrImmune data.
**a)** UMAP projection of the Bcell coloured by (i) sample source, (ii) level of BCR somatic hypermutation (SHM), (iii) BCR isotype usage, and (iv) number of BCR UMIs (in percentiles).
**b)** Correlation between IGH BCR UMIs and IGK/L BCR UMIs.
**c)** Boxplots of the level of (i) BCR SHM and (ii) number of BCR UMIs by B cell subtype.

**d)** Table of the percentage of B cells within each subpopulation expressing each isotype.
**e)** Gene signatures for the detailed B cell subpopulations.

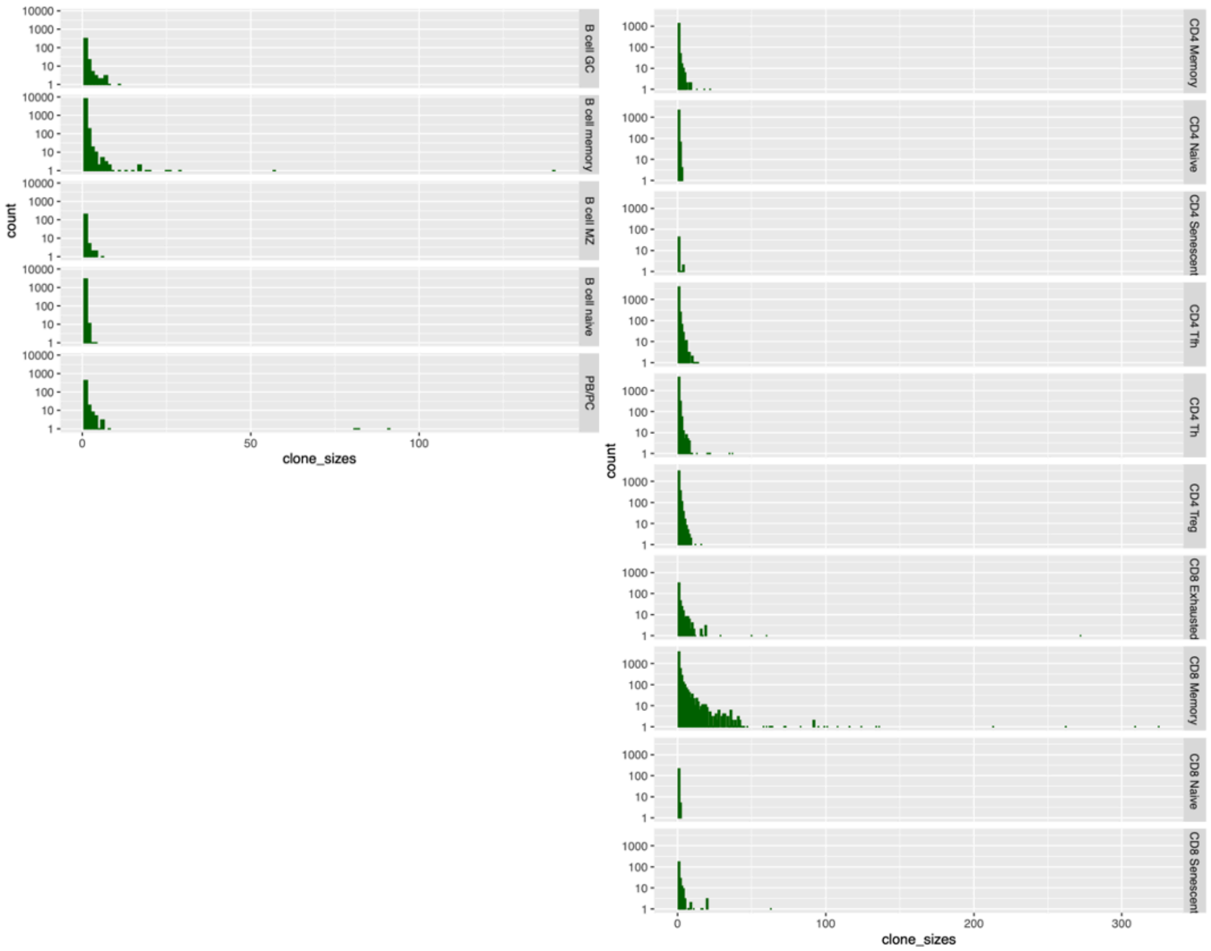


**Supplemental item figure 1.4.** Clone sizes by (left) B cell subpopulation and (right) T cell subpopulation.

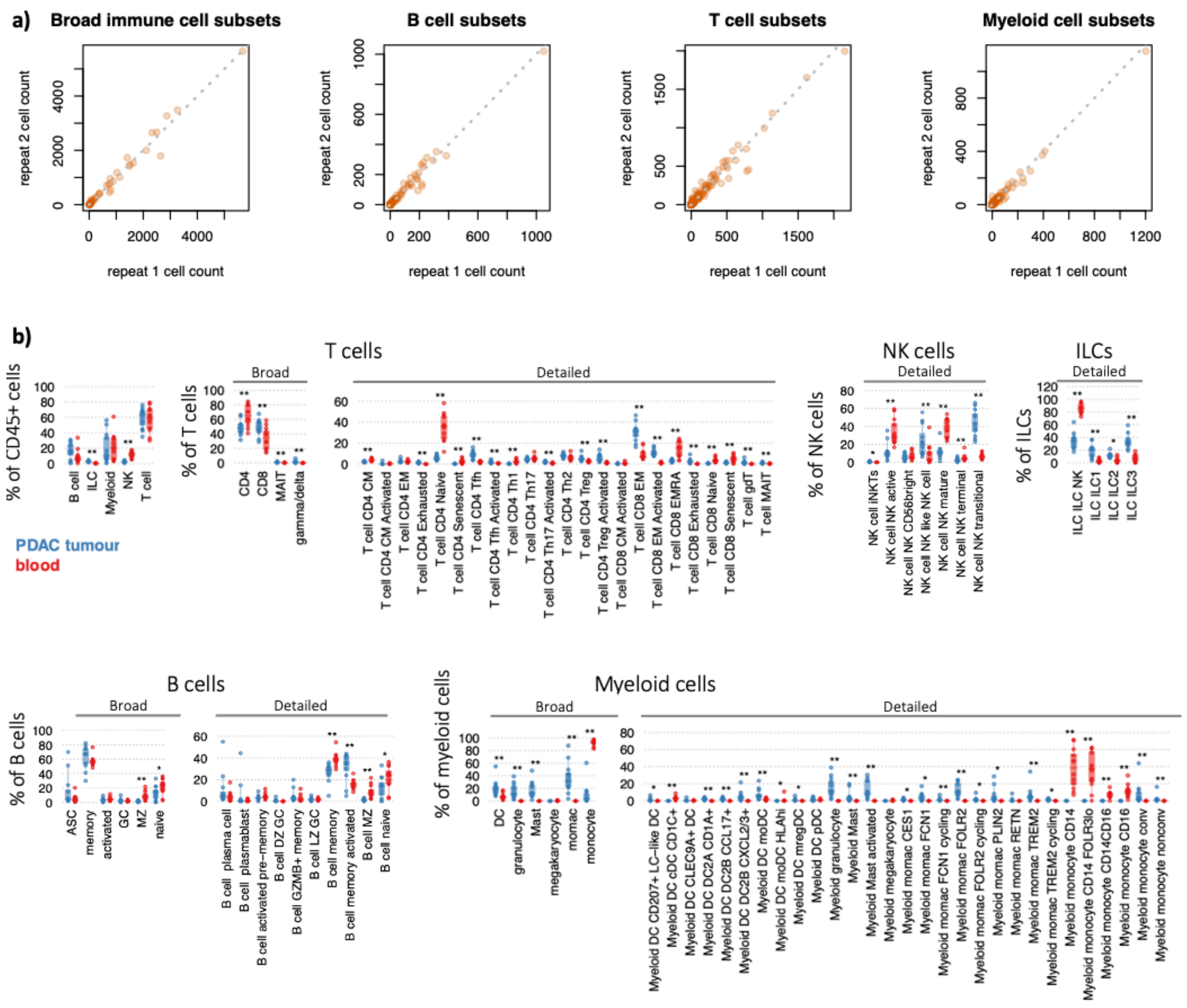


**Supplemental item figure 1.5.** Technical validation and comparisons between blood and tissue.

**a)** Correlations between the cellular subpopulation percentage frequencies between technical repeats, for all cell types and split by broad cell subsets. Technical repeats were dissociated tissue samples that were run on independent 10X channels. The dotted line denotes x=y.

**b)** Immune cell subset proportions between tumour (blue) and blood (red) samples within broad cellular subsets as a proportion within the PancrImmune dataset. * denotes p-values<0.05, ** denotes p-values<0.005 and tests were performed by MANOVA.

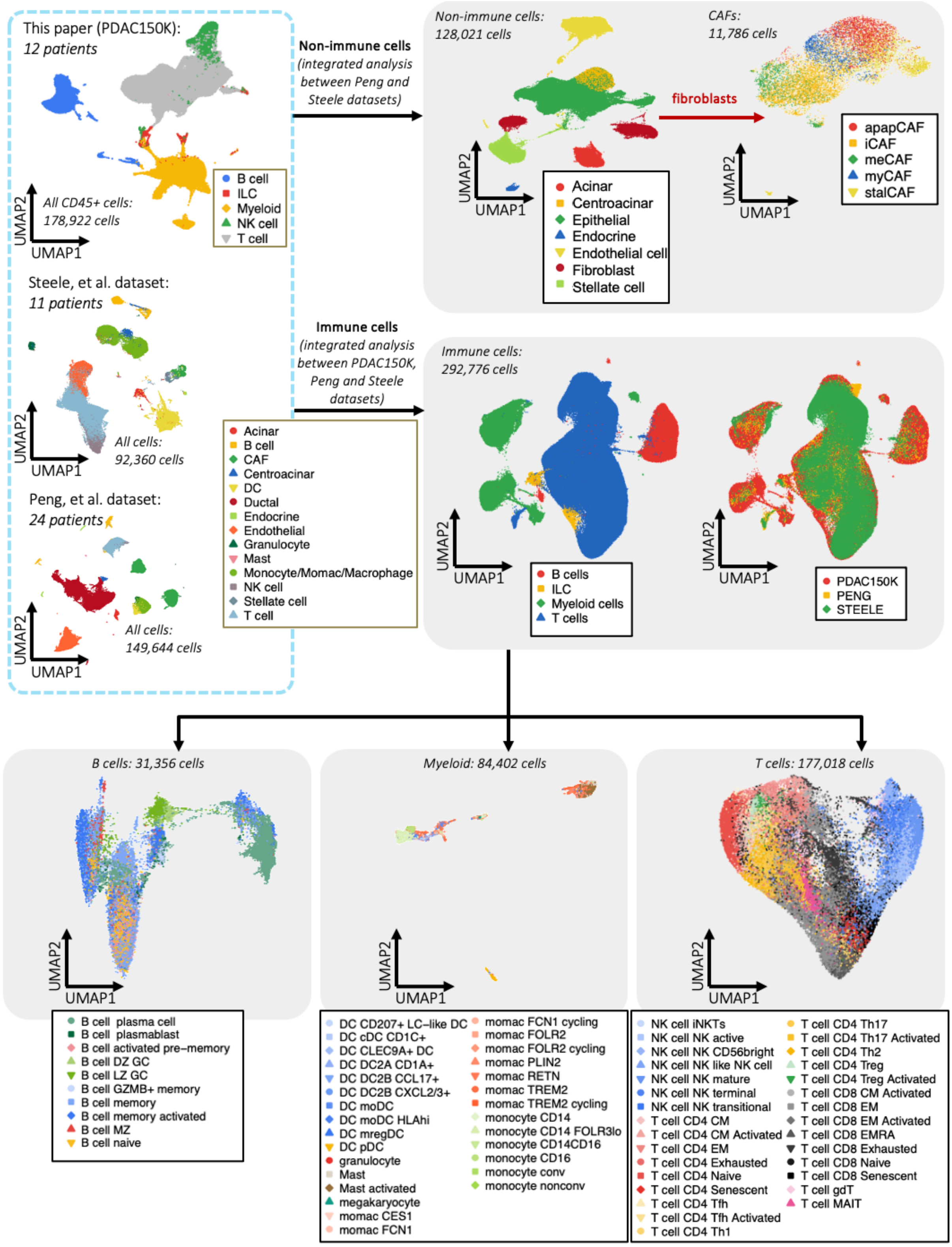


**Supplemental item figure 1.6.** Integration of the PDAC150K, Peng, *et al.* and Steele, *et al.* datasets.

**ai)** Immune cells *(integrated analysis between PDAC150K, Peng and Steele datasets)*

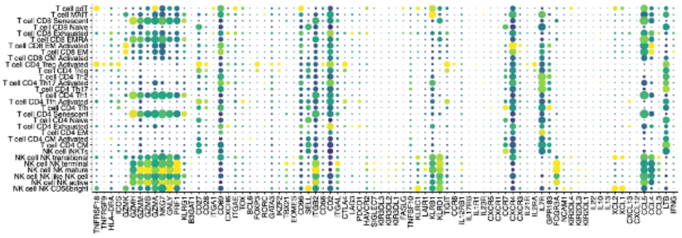

**ii)**

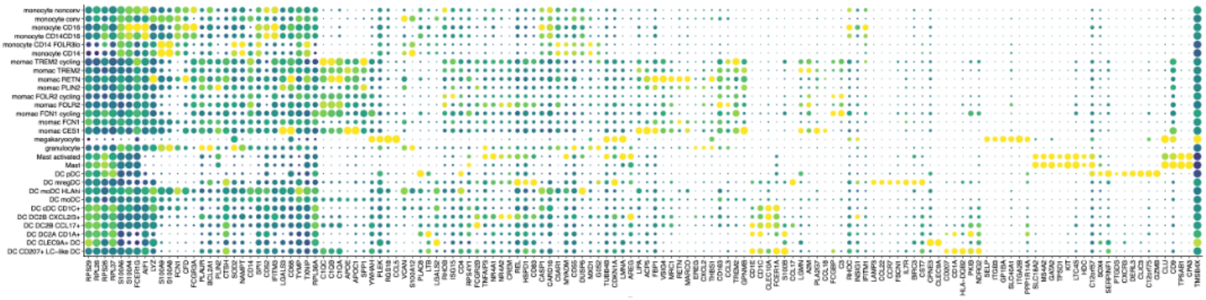

**iii)**

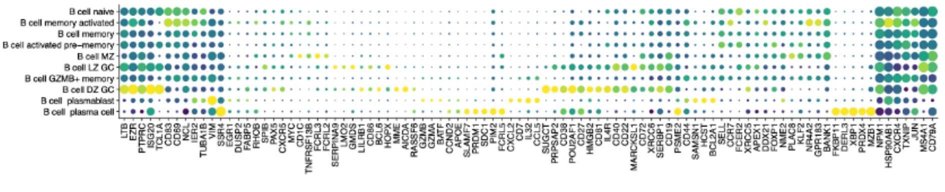


**bi)** Non-immune cells *(integrated analysis between Peng and Steele datasets)*

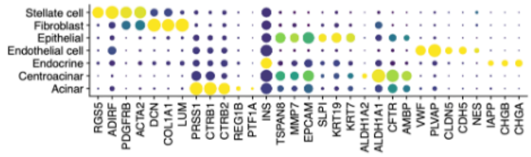


**ii)**

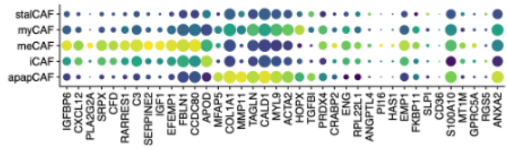


**Supplemental item figure 1.7.** Gene signatures for the cell annotations from the integrated PDAC150K, Peng, *et al.* and Steele, *et al.* datasets for the **a)** immune cell subpopulations and **b)** non-immune cell subpopulations.
